# LncRNA *Dnmt3aos* regulates *Dnmt3a* expression leading to aberrant DNA methylation in macrophage polarization

**DOI:** 10.1101/514307

**Authors:** Xueqin Li, Yingying Zhang, Mengying Zhang, Xiang Kong, Hui Yang, Min Zhong, Weiya Pei, Yang Xu, Xiaolong Zhu, Tianbing Chen, Jingjing Ye, Kun Lv

**Affiliations:** Central Laboratory of Yijishan Hospital, Wannan Medical College, Wuhu, 241001, PR China; Non-coding RNA Research Center of Wannan Medical College, Wuhu, 241001, PR China; Laboratory Medicine of Yijishan Hospital, Wannan Medical College, Wuhu, 241001, PR China

**Keywords:** LncRNA, DNA methylation, Dnmt3a, macrophage polarization

## Abstract

Long non-coding RNAs (lncRNAs) play key roles in various biological processes. However, the roles of lncRNAs in macrophage polarization remain largely unexplored. In this study, thousands of lncRNAs were identified that are differentially expressed in distinct polarized bone marrow-derived macrophages (BMDMs). Among them, *Dnmt3aos* (DNA methyltransferase 3A, opposite strand), as a known lncRNA, locates on the antisense strand of *Dnmt3a*. Functional experiments further confirmed that *Dnmt3aos* were highly expressed in M(IL-4) macrophages and participated in the regulation of *Dnmt3a* expression, and played a key role in macrophage polarization. The DNA methylation profiles between the *Dnmt3aos* knockdown group and the control group in M(IL-4) macrophages were determined by MeDIP-seq technique for the first time, and the *Dnmt3aos-Dnmt3a* axis-mediated DNA methylation modification-regulated macrophage polarization related gene *IFN-γ* was identified. Our study will help to enrich our knowledge of the mechanism of macrophage polarization and will provide new insights for immunotherapy in macrophage-associated diseases.

## Introduction

Macrophages, as an essential component of the innate immune system, are widely distributed throughout the organs and tissues and possess high heterogeneity and plasticity. Macrophages can be broadly classified into two extremely polarized phenotypes, “Classical” and “Alternative,” which exhibit different functions under the influence of distinct microenvironmental stimuli. Classically activated macrophages have long been known to be induced by lipopolysaccharides (LPS) and interferon-gamma (IFN-γ) or granulocyte-macrophage colony stimulating factor (GM-CSF), and are characterized by high levels of pro-inflammatory cytokines such as interleukin-12 (IL-12) and tumor necrosis factor alpha (TNF-α) and oxidative metabolites such as nitric oxide (NO). M(LPS+IFN-γ) macrophages generally promote Th1-type immune responses, with upregulation of MHC II and the costimulatory molecules CD80 and CD86, which are essential for antigen presentation and which strengthen phagocytosis and killing of pathogenic microorganisms or tumor cells [1,2]. In contrast, the alternative M(IL-4) macrophages are induced by Th2 cytokines (such as IL-4 and IL-13) or macrophage colony stimulating factors (M-CSF), with high secretion levels of anti-inflammatory cytokines such as IL-10 and transforming growth factor beta-1 (TGF-β), and high expression of Arginase 1 (Arg-1), chitinase 3-like 3 (Chi3l3 or YM1), and Resistin-like-α (Retnl α or Fizz1). These macrophages participate in tissue repair, angiogenesis, tumor progression, and defense against parasite infection [1,3,4]. Therefore, macrophage polarization is a vital component in the progression of many diseases, including infection [5], tumors [2,6], obesity, and insulin resistance [7,8]. Further study of the molecular mechanisms regulating macrophage polarization will enhance our understanding of the pathogenesis and development of macrophage-centered diseases and provide a basis for the diagnosis and treatment of such diseases. However, the regulatory mechanism of the macrophage activation conditioning response remains to be further elucidated.

LncRNAs are RNA molecules > 200 nucleotides long. They are therefore structurally similar to mRNA, but they do not encode proteins. The lncRNAs are emerging as important regulators of gene transcriptional regulation, chromatin remodeling, nuclear transportation, and other biological processes [9]. They play pivotal roles in various human diseases. Recent studies have confirmed that lncRNAs correlate with macrophage differentiation and activation [10–14]. Epigenetic regulation, including DNA methylation, is critical in cell differentiation and development. *De novo* DNA methylation is catalyzed by DNA methyltransferase (DNMT) 3a and 3b. DNA methylation of cytosines at CpG dinucleotides is the most common epigenetic modification, and CpGs are often enriched in the promoter region of genes [15]. Promoters of hypo-methylated genes are typically associated with transcriptionally active genes [16], whereas DNA hyper-methylation can result in gene silencing [17,18]. However, there are few reports on the role of epigenetic modification in macrophage polarization so far. Recently, histone methylation and Dnmt3b mediated DNA methylation have been implicated in the regulation of macrophage polarization [15,19–21]. However, at present, no studies have been reported on the lncRNAs regulating Dnmt3a expression-mediated aberrant methylation of DNA in macrophage polarization.

In the current study, we utilized microarray and bioinformatics methods to acquire differentially expressed lncRNAs profiles existing in M(LPS+IFN-γ) and M(IL-4) macrophages, and we identify novel lncRNAs, which are functional in macrophage polarization. Among the thousands of lncRNAs that are actively regulated in macrophage polarization, we further determined that lncRNA *Dnmt3aos*, a known lncRNA locates on the *Dnmt3a* antisense strand, is a key regulator of *Dnmt3a* expression and also regulates macrophage polarization. We also used a methylated DNA immunoprecipitation sequencing (MeDIP-seq) method combined with mRNA expression profile analysis to screen the *Dnmt3aos-Dnmt3a* axis-mediated DNA methylation-modified macrophage polarization related genes, and we explored the roles of related genes in macrophage polarization. Our study will help to enrich our understanding of the mechanism of macrophage polarization and will provide new approaches to immunotherapy in macrophage-centered diseases.

## Results

### Identification of *ex vivo* polarized M(LPS+IFN-γ) and M(IL-4) macrophage LncRNAs, including *Dnmt3aos*

To use a systematic approach to identifying alterations in the lncRNA expression profile during macrophage polarization, we examined the lncRNA and mRNA expression profiles in polarized BMDMs through microarray analysis. First, we prepared the M(LPS+IFN-γ) and M(IL-4) macrophages *in vitro*. BMDMs were isolated from BALB/c mice and stimulated with LPS and IFN-γ to obtain M(LPS+IFN-γ) or with IL-4 to obtain M(IL-4). The polarization conditions used in this study resulted in distinct macrophage phenotypes as confirmed in our previous study [22–24].

For the microarray analysis, authoritative data sources containing more than 33,231 lncRNAs were used. To identify the most significant candidates, lncRNAs with at least two-fold differentially expressed changes and with FDR adjusted *P* values < 0.05 were selected (Fig 1a and 1b and S1 Data). Under these criteria, 627 lncRNAs were up-regulated, and 624 lncRNAs were down-regulated in the M(LPS+IFN-γ) group compared with the M(IL-4) group. In addition, 696 mRNAs were up-regulated and 557 mRNAs were down-regulated in the M(LPS+IFN-γ) group compared with the M(IL-4) group (Tab 1 and S1 Data). The mRNA expression profiles were reported in our previous publication [22].

**Fig 1.**
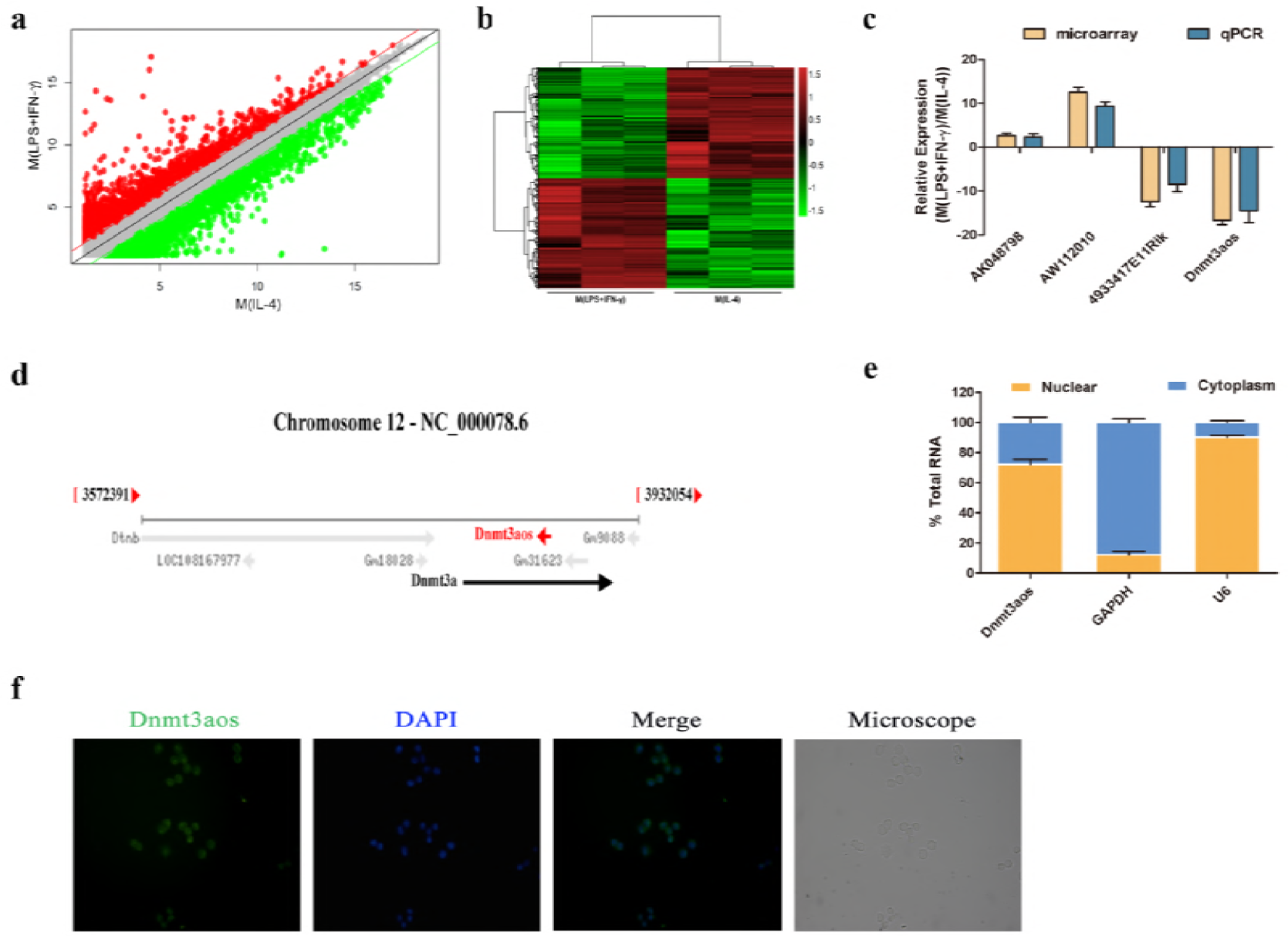
Identification and characterization of *Dnmt3aos*. **(a-b)** LncRNA microarray expression data from BMDMs incubated in distinct polarizing conditions. A total of 1,251 differentially-expressed lncRNAs were identified between the M(LPS+IFN-γ) and M(IL-4) macrophages. **(a)** Scatter plots show the variation in lncRNA expression between the M(LPS+IFN-γ) and M(IL-4) macrophages. The values of the X and Y axes in the scatter plot are the averaged normalized values in each group (log2-scaled). **(b)** Heat maps of lncRNAs expression profiles between the two groups. “Red” indicates high relative expression and “green” indicates low relative expression. One ANOVA test was used for statistical analysis. LncRNA with expression fold change > 2 and with FDR-adjusted *P* value < 0.05 was considered statistically significant. **(c)** Confirmation of the differential expression of lncRNAs by RT-qPCR. Four differentially expressed lncRNAs were validated by RT-qPCR. The y-axis represents the log2-transformed median fold-change in expression. **(d)** LncRNA Dnmt3aos locates on the Dnmt3a opposite strand on mouse chromosome 12. **(e)** RT-qPCR detection of *Dnmt3aos* level in cellular fractions from BMDMs. U6 and GAPDH were the nuclear and cytoplasmic controls, respectively. Data were expressed as the means ± SD of three independent experiments. **(f)** *Dnmt3aos* expression in BMDMs detected by RNA-FISH. Scale bar, 20 mm.

To confirm the microarray results, four differentially expressed lncRNAs were selected for further confirmation using RT-qPCR (Fig 1c). Among the four selected lncRNAs, *AK048798* and *AW112010* were upregulated in M(LPS+IFN-γ) macrophages compared to M(IL-4) macrophages, while *4933417E11Rik* and *Dnmt3aos* were downregulated in M(LPS+IFN-γ) macrophages compared to M(IL-4) macrophages (Fig 1c). The results were generally consistent with the microarray data.

Since many lncRNAs have been shown to regulate neighboring genes either positively or negatively [25–27] as their target genes, the genomic locations of these lncRNAs were further characterized. We decided to focus on lncRNA *Dnmt3aos* since it is a known lncRNA, locates on the *Dnmt3a* gene antisense strand on mouse chromosome 12 (Fig 1d).

### Characterization of the *Dnmt3aos* expression pattern

To further determine the cellular localization of the *Dnmt3aos* transcript, the nuclear and cytosolic RNAs were isolated from BMDMs, and the expressions of *Dnmt3aos* transcripts in both subcellular locations were measured. RT-qPCR data show that *Dnmt3aos* transcripts are highly expressed in the nucleus compared with the cytosol (Fig 1e). As controls, *GAPDH* mRNA is specifically located in the cytosol, whereas *U6* RNA is primarily located in the nucleus (Fig 1e). RNA fluorescence in situ hybridization (RNA FISH) indicates that *Dnmt3aos* is highly expressed in the nucleus (Fig 1f).

We next examined the expression profile of *Dnmt3aos* in multiple mouse tissues. *Dnmt3aos* was expressed in many mouse tissues (S1 Fig), including lung, brain, spleen, pancreas, liver, heart, kidney, and fat.

### Elevated expression of *Dnmt3aos* in M(IL-4) polarized macrophages contributes to macrophage polarization

*Dnmt3aos* had increased expression in M(IL-4) macrophages (Fig 2a). Therefore, we further characterized its role in macrophage activation. We assessed the level of *Dnmt3aos* in macrophages after the dynamic process of macrophage re-polarization.

**Fig 2.**
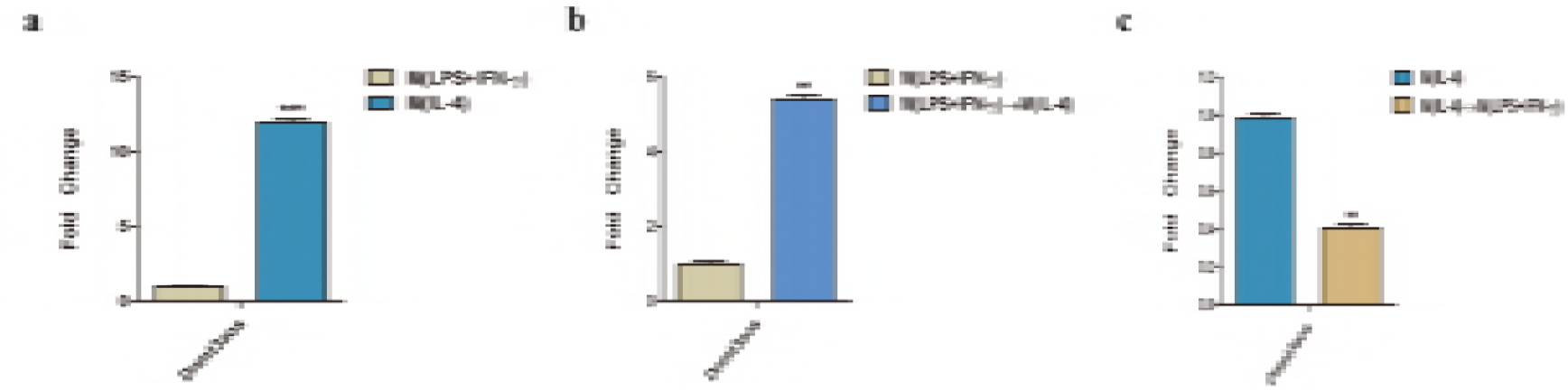
Differential expression of *Dnmt3aos* during macrophage polarization. **(a)** *Dnmt3aos* was assessed by RT-qPCR and normalized to GAPDH in BMDMs after 48 h of stimulation with LPS (100 ng/ml) plus IFN-γ (20 ng/ml) or IL-4 (20 ng/ml). **(b)** *Dnmt3aos* levels in macrophages following M(LPS+IFN-γ)-to-M(IL-4) re-polarization by IL-4 (20 ng/ml) for 24 h. **(c)** *Dnmt3aos* levels in macrophages following M(IL-4)-to-M(LPS+IFN-γ) re-polarization by LPS (100 ng/ml) plus IFN-γ (20 ng/ml) for 8 h. Data are expressed as the means ± SD of three independent experiments. \*\**P* < 0.01, \*\*\**P* < 0.001, *P*-value was calculated based on Student’s *t* test.

Macrophages with the M(LPS+IFN-γ) or M(IL-4) phenotypes were re-polarized to the M(IL-4) or M(LPS+IFN-γ) phenotype by treatment with LPS/IFN-γ or IL-4, respectively. The marker gene (we used iNOS as the marker for M(LPS+IFN-γ) macrophages and Arg1 as the marker for M(IL-4) macrophages [22]) assays clearly showed that M(LPS+IFN-γ) macrophages could be re-polarized to M(IL-4) macrophages by IL-4 (S2a Fig), and M(IL-4) macrophages could be re-polarized to M(LPS+IFN-γ) macrophages by LPS/IFN-γ (S2b Fig). Interestingly, the *Dnmt3aos* levels in macrophages were strikingly elevated during M(LPS+IFN-γ)-to-M(IL-4) re-polarization of macrophages (Fig 2b), but *Dnmt3aos* levels decreased following M(IL-4)-to-M(LPS+IFN-γ) re-polarization of macrophages (Fig 2c). These results suggest that *Dnmt3aos* may play a critical role in promoting macrophage polarization to the M(IL-4) phenotype.

### *Dnmt3aos* regulates *Dnmt3a* RNA and protein in BMDMs

To test the correlation between *Dnmt3aos* levels and macrophage polarization, the gene was examined after primary BMDMs were treated with LPS/IFN-γ or IL-4 stimulation. The *Dnmt3aos* gene was down-regulated with the polarization of M(LPS+IFN-γ) macrophages, but up-regulated with the M(IL-4) polarization (Fig 3a) when compared to the primary BMDMs. Meanwhile, the expression level followed a time course dependent on the LPS/IFN-γ or IL-4 stimulation (Fig 3a), and correlated to the M(LPS+IFN-γ) and M(IL-4) phenotype, marked by iNOS and Arg1 gene expression, respectively (S3 Fig). Notably, this expression pattern was almost identical to that of Dnmt3a mRNA (Fig 3b). This observation suggests that *Dnmt3aos* and *Dnmt3a* expression may be regulated concordantly, as was recently shown for other sense-antisense pairs [28–31],

**Fig 3.**
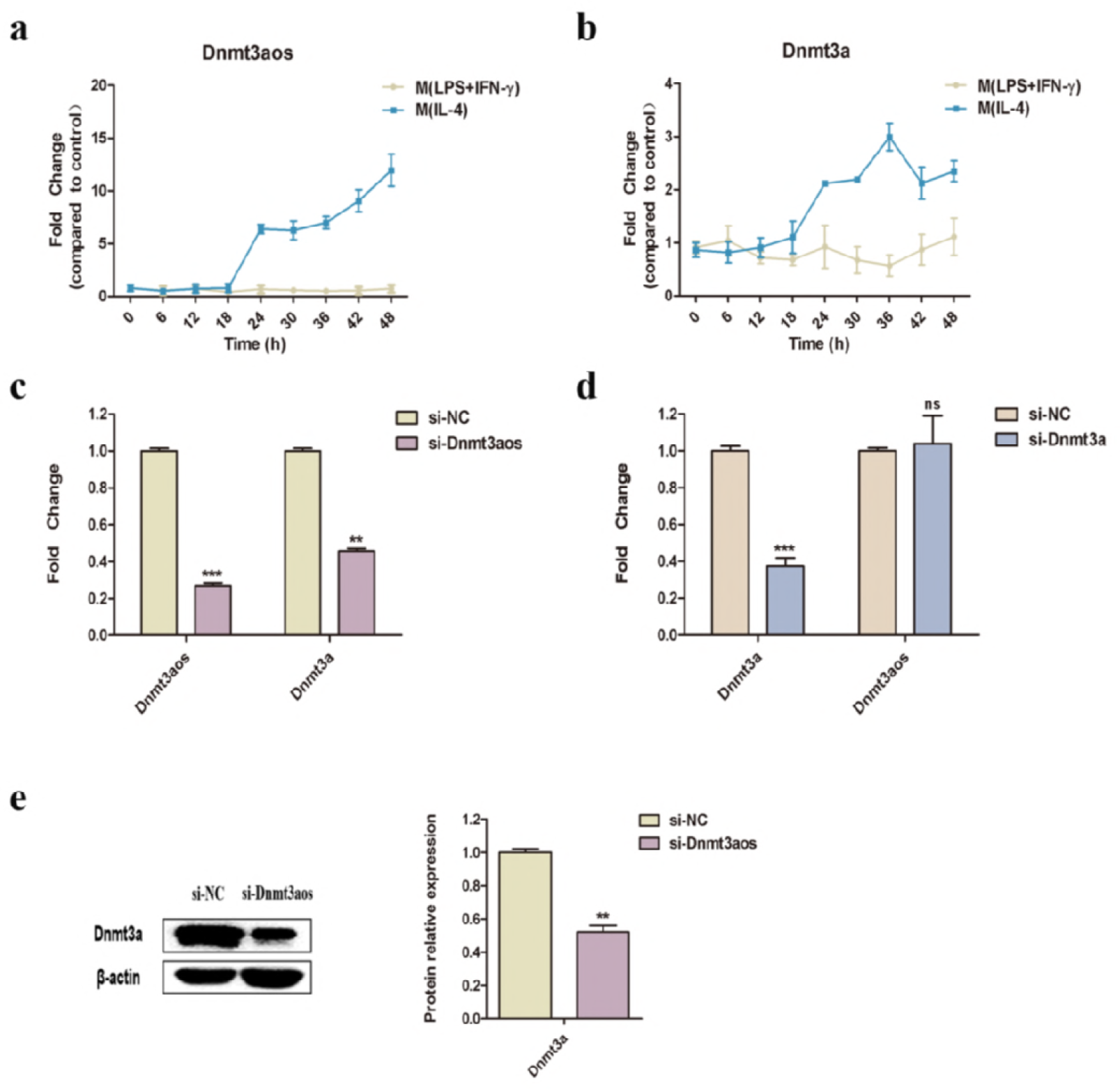
*Dnmt3aos* regulates *Dnmt3a* mRNA and protein expression *in vitro*. **(a-b)** Time course of *Dnmt3aos* **(a)** or *Dnmt3a* **(b)** expression in BMDMs after stimulation with LPS (100 ng/ml) plus IFN-γ (20 ng/ml) or IL-4 (20 ng/ml). **(c-d)** Knockdown of *Dnmt3aos* **(c)** or *Dnmt3a* **(d)** in BMDMs with lncRNA smart silencer (si-Dnmt3aos) or siRNA specifically targeting Dnmt3a (si-Dnmt3a). *Dnmt3aos* and *Dnmt3a* expression were assessed by RT-qPCR. Nontargeting siRNA negative control (si-NC) was used as the control. **(e)** Knockdown of *Dnmt3aos* in BMDMs with si-Dnmt3aos or si-NC. Protein expression of *Dnmt3a* and *β-actin* set as the endogenous control were measured by western blotting. Data are expressed as the means ± SD of two independent experiments. \*\**P* < 0.01, \*\*\**P* < 0.001. *P*-value was calculated based on Student’s *t* test.

We next investigated whether the *Dnmt3aos* transcript regulates expression of *Dnmt3a* mRNA. RT-qPCR results showed that transfection of primary BMDM cells with Dnmt3aos smart silencer (si-Dnmt3aos) resulted in a statistically significant knockdown of not only the targeted *Dnmt3aos* transcript, but also of *Dnmt3a* mRNA and protein levels (Fig 3c and 3e). In contrast, the expression of the si-Dnmt3a in primary BMDMs resulted in a significant reduction of the targeted *Dnmt3a* mRNA and protein levels (Fig 3d and S4 Fig), but did not affect the expression of *Dnmt3aos* transcripts (Fig 3d). This suggests that *Dnmt3aos* may regulate the expression of *Dnmt3a* at both the mRNA and the protein levels in primary BMDMs. These observations further confirm the regulation of *Dnmt3a* expression by *Dnmt3aos*, not only at the mRNA but also at the protein level.

### Knockdown of *Dnmt3aos* expression promotes M(LPS+IFN-γ) macrophage polarization and decreases M(IL-4) macrophage polarization

The expression profile of *Dnmt3aos* prompted us to investigate its function during macrophage polarization. To determine if *Dnmt3aos* participates in macrophage polarization, we transfected primary BMDM macrophages with Dnmt3aos smart silencer. Interestingly, when primary BMDM macrophages were transfected with Dnmt3aos smart silencer and then treated with LPS/IFN-γ or IL-4, the M(LPS+IFN-γ) phenotype was increased. Specifically, the M(LPS+IFN-γ) macrophage phenotype markers iNOS, TNF-α, and IL-12 were all up-regulated, but the M(IL-4) macrophage markers Arg1, YM1, and FIZZ1 were strikingly down-regulated (Fig 4). These results suggested that *Dnmt3aos* may play a critical role in promoting macrophage polarization.

**Fig 4.**
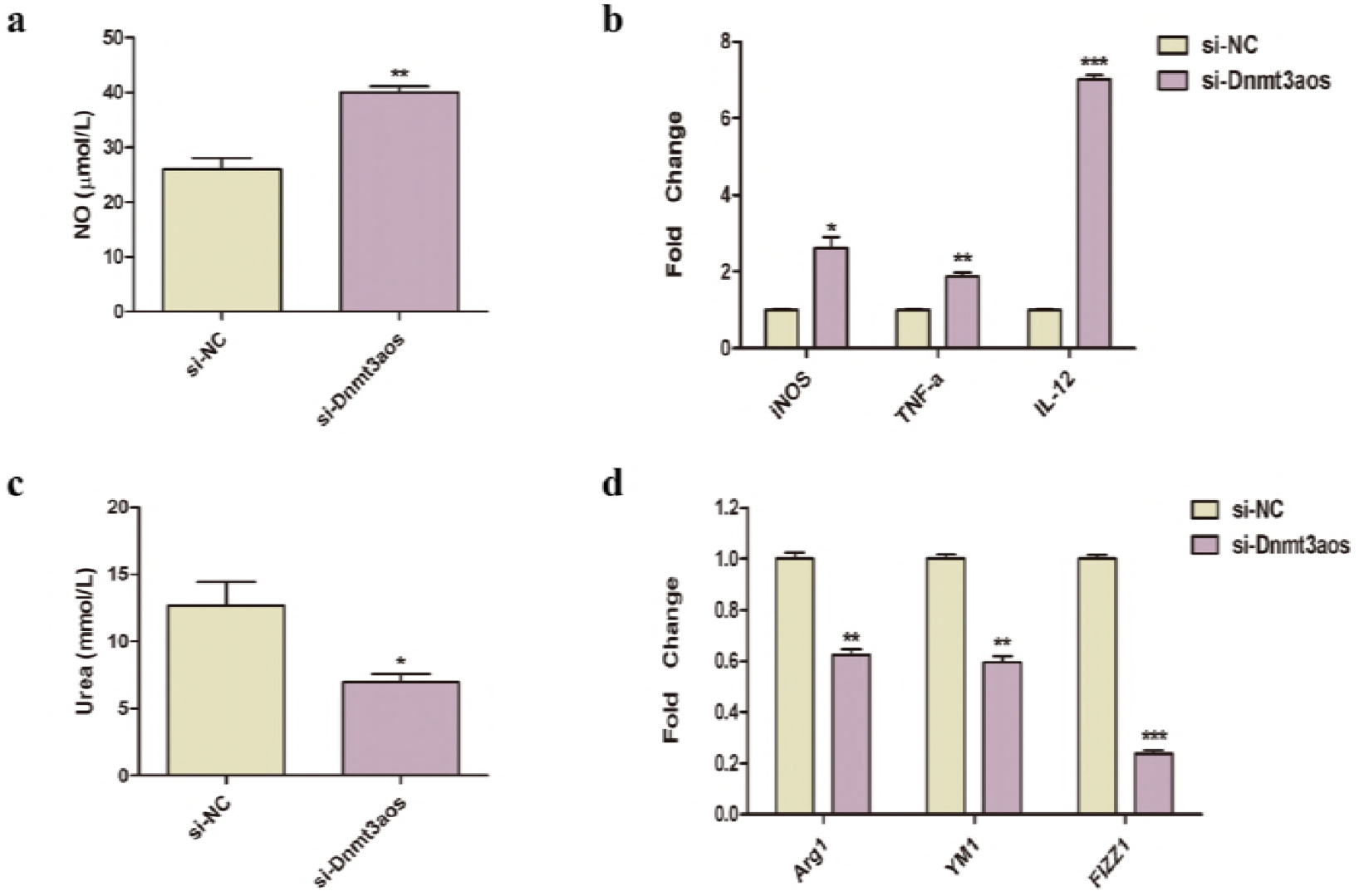
*Dnmt3aos* plays important roles in BMDMs polarization. **(a-b)** Primary BMDMs were transfected with si-NC or si-Dnmt3aos for 48 h, followed by stimulation with LPS (100 ng/mL) plus IFN-γ (20 ng/ml) for an additional 6 h. **(a)** Supernatants from the two groups of cells were observed for NO production using Griess activity assays. **(b)** RT-qPCR. RNA was extracted from the two groups of cells and subjected to RT-qPCR analysis with primers specific to iNOS, TNF-α, and IL-12. **(c-d)** Primary BMDMs were transfected with si-NC or si-Dnmt3aos for 48 h, followed by stimulation with IL-4 (20 ng/ml) for an additional 24 h. **(c)** Cells were lysed and investigated for urea production using arginase activity assays. **(d)** RT-qPCR. RNA was extracted from the two groups of cells and subjected to RT-qPCR analysis with primers specific to Arg1, YM1, and FIZZ1. GAPDH was set as the endogenous control. Data are expressed as the means ± SD of three independent experiments. \**P* < 0.05, \*\**P* < 0.01, \*\*\**P* < 0.001. *P*-value was calculated based on Student’s *t* test.

### The *Dnmt3aos-Dnmt3a* axis regulates macrophage polarization-related gene expression through modification of DNA methylation

The establishment and maintenance of *de novo* DNA methyltransferase 3A (Dnmt3a) mediated methylation patterns resulting in modulation of gene expression is one of the key steps in epigenetic regulation during cell differentiation and biological processes [32–35]. Since the *Dnmt3aos* and *Dnmt3a* genes were differentially expressed in macrophage polarization and were highly expressed in M(IL-4) macrophages (Fig 3a and 3b), and based on the results that *Dnmt3aos* regulates *Dnmt3a* gene expression, we carried out methylated DNA immunoprecipitation sequencing (MeDIP-seq) to evaluate the possibility of association between DNA methylation variability and macrophage polarization to identify differentially methylated regions (DMRs) related to macrophage polarization. Indeed, unique DNA methylation features were identified in negative control non-targeting siRNA- (si-NC-) and si-Dnmt3aos-transfected M(IL-4) macrophages. We looked at methylation changes in four regions defined by the distance from the CpG islands [36], namely, the CpG island, Shore, Shelf, and Open Sea regions, the latter three being, respectively, 2 kb, 2-4 kb, and more than 4 kb from the CpG island. Most CG islands of the si-NC group and si-Dnmt3aos group both were detected in the Open Sea regions (Fig 5a). Next, we scrutinized the distribution of DMRs in the context of gene structure. Seven relevant regions were defined: promoter, five-prime untranslated region (UTR), coding sequence (CDS), three-prime UTR, intron, the transcriptional termination region (TTR), and the intergenic region. However, the DMRs were distributed differentially in the seven regions, and had no significant changes between the si-NC group and si-Dnmt3aos group. As shown in Fig 5b, DMRs were concentrated mainly in introns and in the intergenic region. Heat maps of DMRs methylation levels for the two biological replicates of the si-NC group and si-Dnmt3aos group are shown in Fig 5c. In total, 5,143 DMRs were selected based on the criterion of fold change (FC) >1.5 and corrected *P* value (FDR) < 0.05 (S2 Data). The chromosomal locations of the DMRs are shown in Fig 5d. The red represents high methylation and the blue represents low methylation. The height represents logFC. Circos plots of global genomic methylation level across each chromosome with comparisons between the si-NC group and si-Dnmt3aos group are shown in Fig 5e. On the whole, the chromosomal locations between the si-NC and si-Dnmt3aos-transfected M(IL-4) macrophages were almost identical.

**Fig 5.**
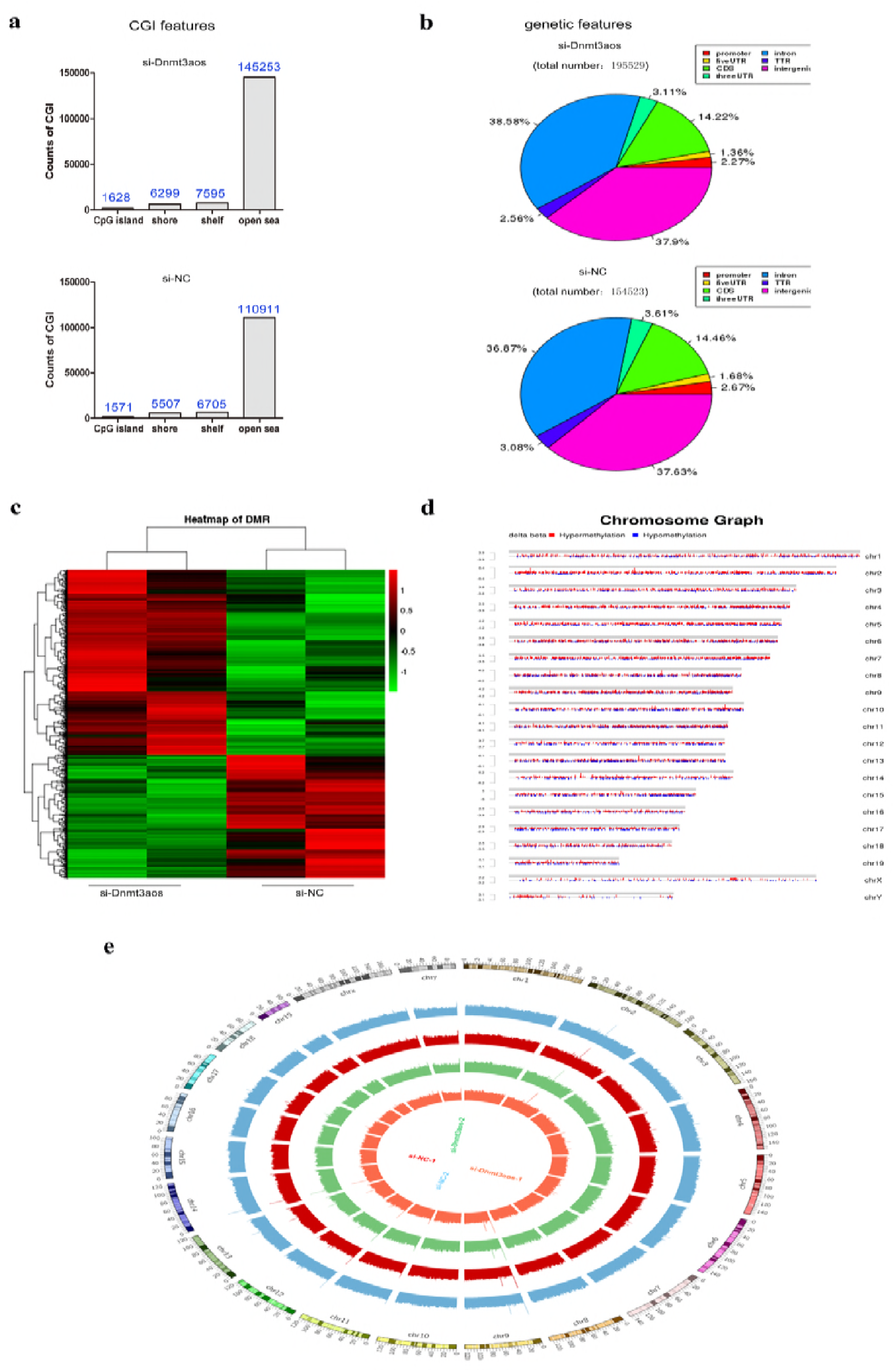
MeDIP-seq methylation profiles between the si-NC and si-Dnmt3aos transfected M(IL-4) macrophages. **(a)** Distribution and counts of CpG islands (CGI) obtained from the si-NC and si-Dnmt3aos transfected M(IL-4) macrophages are shown in the histogram. **(b)** The average proportions of CGI obtained from the si-NC and si-Dnmt3aos-transfected M(IL-4) macrophages in each region defined by genomic structure are shown in pie charts. Promoters (−2 kb regions around their TSSs), CpG islands and their Shores (up/down stream 2 kb from the CpG island), Shelf (up/down stream 2-4 kb from the CpG island), and Open Sea (more than 4 kb from the CpG island). **(c)** Heat map of differentially methylated regions (DMRs), selected by highest variances between samples, including mean rpm signals for the two biological replicates of the si-NC and si-Dnmt3aos-transfected M(IL-4) macrophages. Red indicates up-regulated fold change and green indicates down-regulated fold change. **(d)** Chromosomal distribution of DMRs. Red represents high methylation and blue represents low methylation. The height represents logFC (fold change). **(e)** Circos plot of global genomic methylation level across each chromosome between the si-NC and si-Dnmt3aos-transfected M(IL-4) macrophages. The coverage of the genome is observed by a 100 K window, indicating the methylation level of the genome; the outermost circle in the figure is the chromosomes, and each circle inside indicates the genome methylation level of the sample.

To generate insights into the potential biological functions of DMRs, functional enrichment analysis was performed using GO and KEGG pathway terms. Based on DMR-related genes, we provided a GO functional classification annotation for DMR-related genes. All differently expressed genes were mapped to each term of the GO database (http://www.geneontology.org/) and the KEGG pathway database (https://www.kegg.jp/kegg/pathway.html), the gene numbers of the GO and KEGG terms were calculated. A hypergeometric test was used to determine significantly enriched GO and KEGG terms of the DMR-related genes compared to the genome background, and the corrected-*P* value ≤ 0.05 was used as a threshold. GO and KEGG terms fulfilling this condition were defined as significantly enriched GO and KEGG terms of DMR-related genes. Through this analysis, we were able to recognize the top 30 biological functions (Fig 6a and S3 Data) and top 30 pathways in which DMR-related genes participated (Fig 6b and S4 Data).

**Fig 6.**
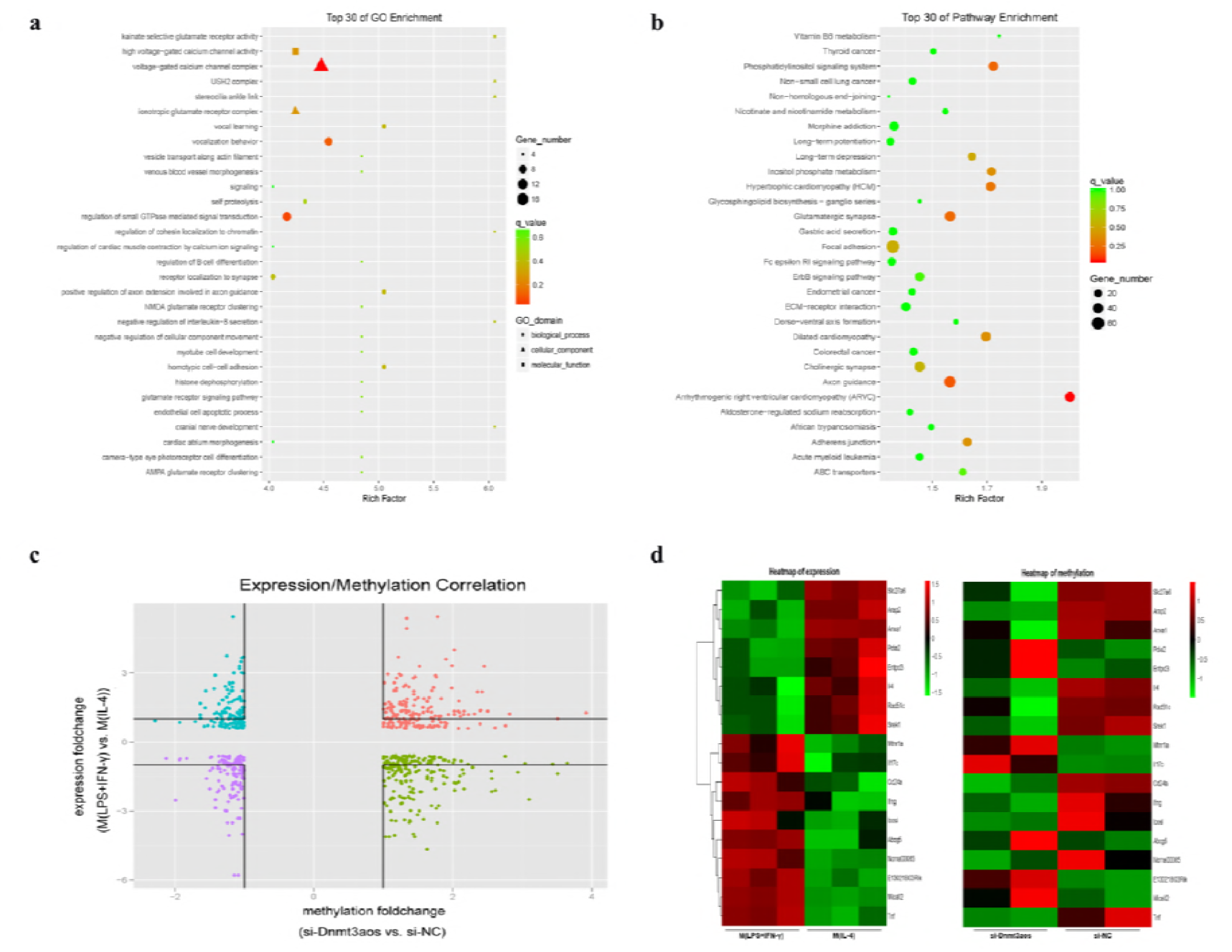
Enrichment analysis of DMRs. **(a-b)** Analysis of the top 30 GO enrichment terms **(a)** and the top 30 KEGG pathway enrichment terms **(b)** of DMRs. For each term, the expected and observed gene numbers along with the statistical significance (q-value) for the enrichments are presented. **(c-d)** Correlation analysis of differentially methylated region (DMR) methylation status of the si-NC and si-Dnmt3aos-transfected M(IL-4) macrophages and differentially expressed genes (DEGs) of M(LPS+IFN-γ) and M(IL-4) macrophages. **(c)** Overall view of the negative correlation between methylation and expression. DEGs are shown on the longitudinal coordinates to show the difference in expression, and the difference in methylation is shown on the transverse coordinates. **(d)** Heat map shows the level and correlation of methylation and expression of each gene. The same genes were placed on the same lines.

Our previous investigations of gene expression in macrophage polarization have shown that 1,253 unique genes are significantly differentially expressed genes (DEGs) [22]. Since we have previously confirmed that *Dnmt3aos* plays a critical role in macrophage polarization, knockdown of Dnmt3aos expression promotes the M(LPS+IFN-γ) macrophage phenotype and decreases the M(IL-4) macrophage phenotype (Fig 4). In order to explore the relationship between DNA methylation and gene expression in macrophage polarization, we combined the DNA methylation profiles from the si-NC and si-Dnmt3aos-transfected M(IL-4) macrophages and gene expression profiles (Data are presented in reference 22) to identify the DMR-related genes. As expected, hypomethylated DMRs were associated with increased gene expression (exhibited as blue dots in Fig 6c), and hypermethylated DMRs were associated with decreased gene expression (exhibited as green dots in Fig 6c). Others in which methylation changes were not associated with expected gene expression differences are exhibited as pink and purple dots in Fig 6c and are listed in S5 Data. It is known that the high level of DNA methylation in promoter region inhibits the expression of its genes. Therefore, we choose DMR-related genes which are negatively correlated with the level of DNA methylation in promoter region and its expression level of mRNA for follow-up study. Heat maps were used to show the list of DMR-related genes distributed in promoter regions (Fig 6d). Among these genes, *Ifng, Icosl, Cd24a, Ncrna00085*, and *Tnf* genes were hypermethylated in the si-NC group compared to the si-Dnmt3aos group. These genes were associated with increased expression in M(LPS+IFN-γ) macrophages compared to M(IL-4) macrophages. The *Pdia2* and *Entpd3* genes were hypomethylated in the si-NC compared to the si-Dnmt3aos group. These genes were associated with decreased expression in M(LPS+IFN-γ) macrophages compared to M(IL-4) macrophages (see S6 Data).

Given the finding that the differential expression of *Dnmt3a* in polarized macrophages was dependent on *Dnmt3aos* and was responsible for the DNA methylation of DMR-related genes, we sought to determine whether the expression of DMR-related genes was also regulated by the *Dnmt3aos-Dnmt3a* axis. As expected, the results of this experiment showed that compared to the si-NC group, knockdown of *Dnmt3aos* expression not only down-regulated the M(IL-4) macrophage phenotype related genes (Arg1, YM1, and FIZZ1), but also influenced the expression of the selected DMR-related genes like *IFN-γ, Icosl, Cd24a, TNF-α, Lrrk2, Lox, Pdia2 and Entpd3* (Fig 7a). Additionally, *IFN-γ, Icosl, Cd24a, TNF-α, Lrrk2*, and *Lox* were also elevated in macrophages treated with specific siRNA targeting *Dnmt3a* for knockdown in M(IL-4) (Fig 7b). The same results were observed in M(IL-4) macrophages treated with the DNMT inhibitor 5-Aza-2’-deoxycytidine (5-Aza-CdR) (Fig 7c).

**Fig 7.**
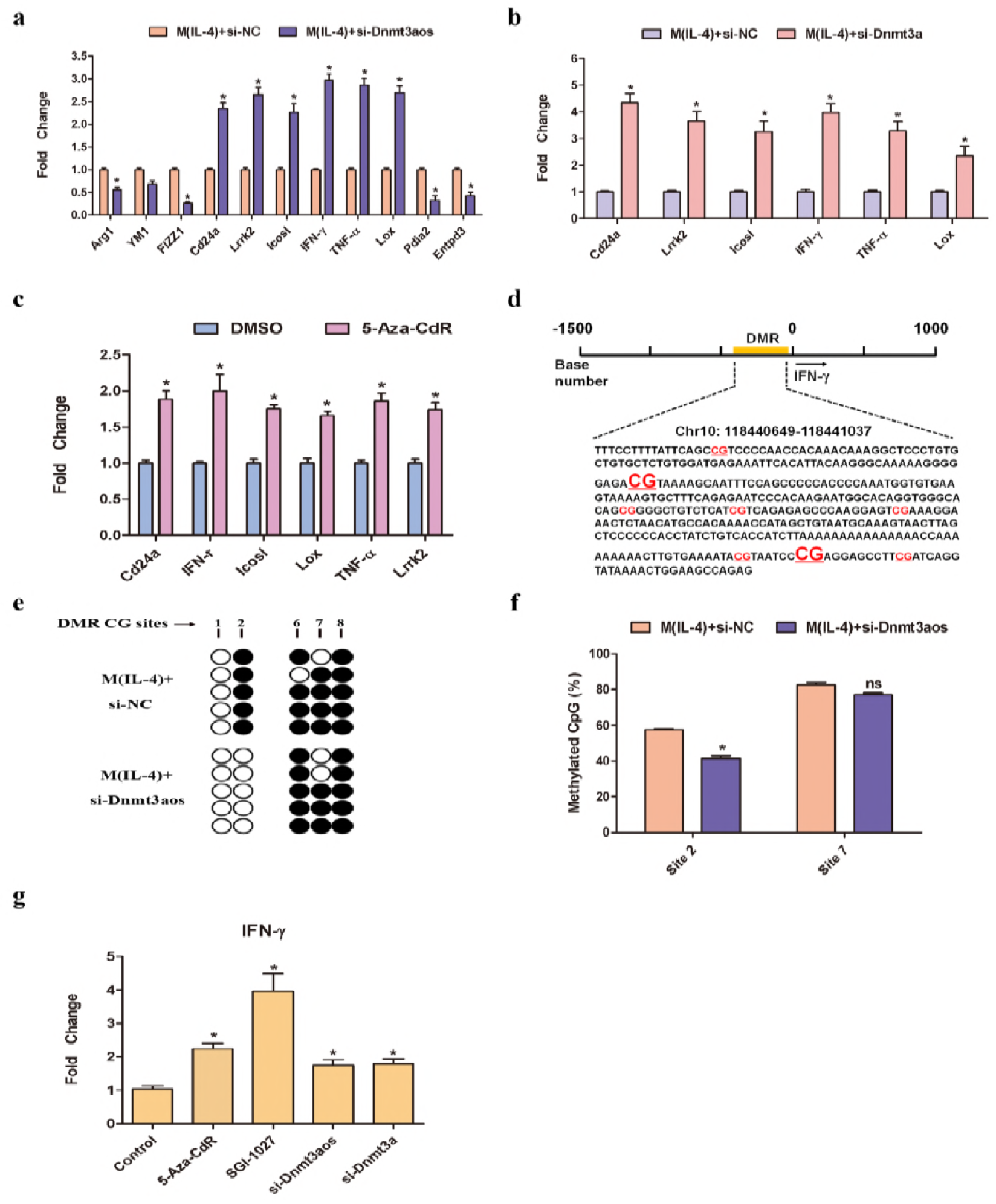
The *Dnmt3aos-Dnmt3a* axis regulates macrophage polarization-related gene expression through modification of DNA methylation. **(a-b)** RT-qPCR validation. Knockdown of *Dnmt3aos* **(a)** or *Dnmt3a* **(b)** expression in M(IL-4) macrophages with lncRNA smart silencer (si-Dnmt3aos) or siRNA specifically targeting Dnmt3a (si-Dnmt3a), respectively. M(IL-4) macrophage phenotype related genes and the selected DMR-related genes obtained from negative correlation analysis of DMRs located in promoter regions and DEGs. **(c)** M(IL-4) macrophages were treated with the DNMT inhibitor 5-Aza-CdR for 72 h. The expression of the selected DMRs (obtained from negative correlation analysis of DMRs located in promoter regions and DEGs) were assessed by RT-qPCR. **(d)** The location and exhibition of DNA sequences of DMRs in the promoter region of *IFN-γ* obtained from MeDIP-seq enrichment. The CG sites (sites 1-2 and sites 6-8 in order) with underlining and red color were successfully BSP sequenced. **(e-f)** *IFN-γ* methylation profiles in the si-NC or si-Dnmt3aos-transfected M(IL-4) macrophages. **(e)** The open and filled circles symbolize the unmethylated and methylated CGs, respectively. Five colonies of separate CG sites from each group were further analyzed by BSP sequencing. **(f)** Pyrosequencing analysis of the CG methylation level in the selected CG sites of the *IFN-γ* gene promoter from BSP sequencing results. The figure illustrates the variation in methylation level at CG site 2 and site 7 in the si-NC or si-Dnmt3aos-transfected M(IL-4) macrophages. **(g)** The expression levels of *IFN-γ* mRNA in M(IL-4) macrophages treated with 5-Aza-CdR, SGI-1027, Dnmt3aos lncRNA smart silencer (si-Dnmt3aos), or Dnmt3a siRNA (si-Dnmt3a). Data were expressed as the means ± SD of three independent experiments. \**P* < 0.05. *P*-value was calculated based on Student’s *t* test.

To further validate the differential methylated CG sites in the DMR region from the MeDIP-seq results (S7 Data), genomic DNA samples of the si-NC and si-Dnmt3aos-transfected M(IL-4) macrophages were treated with sodium bisulfate and amplified by PCR with specific primers. After TA cloning, five positive clones were selected for DNA sequencing. The methylation patterns of these genes are displayed in white and black rectangles (Fig 7e), where the black and white rectangles represent methylated CG and non-methylated CG loci, respectively. The bisulfite sequencing PCR (BSP) sequencing results were analyzed and are summarized in Fig 7e and S5-8 Fig. The whole *IFN-γ* DMR region from the MeDIP-seq results contained 8 CG sites (Fig 7d), but the BSP sequencing successfully sequenced sites 1-2 and 6-8 in order. The white-black rectangle diagram shows that only the site 2 and site 7 CG sites were hypermethylated in the si-NC group compared to the si-Dnmt3aos group and were in accordance with the MeDIP-seq results (Fig 7e). Bisulfite pyrosequencing was then performed to further validate the global DNA methylation percentages of the single CG dinucleotide locus at the site 2 and site 7 CG sites in the *IFN-γ* DMR region, which were differentially methylated according to the BSP results in the si-NC and si-Dnmt3aos-transfected M(IL-4) macrophages. The results showed that the site 2 CG dinucleotide loci of *IFN-γ* CG sites had significant differences between the two group macrophages, while the site 7 CG dinucleotide loci of *IFN-γ* CG sites were calculated as having no difference (Fig 7f). The expression of *IFN-γ* was induced when its promoter was demethylated, and this induction was achieved by Dnmt3a and *Dnmt3aos* knockdown (Fig 7g). These results collectively suggested that *Dnmt3aos-Dnmt3a* axis mediated DNA methylation modification was involved in macrophage polarization.

## Discussion

Macrophage polarization is associated with a variety of diseases, such as infection, tumors, and obesity. Based on the polarization of macrophages into M1 and M2, two extreme categories under different environmental stimuli, the functions of macrophages are completely different and are even mutually inhibited. Appropriate activation of macrophages helps to clear pathogens and tumors, while inappropriate activation of macrophages may inhibit an organism’s immune system, promoting tumor occurrence or progression and chronic infection [37–40]. Macrophage polarization is regulated by different mechanisms, including intracellular signaling pathways, transcription factors, and epigenetic and post-translational modifications. Recent studies have shown that non-coding RNA (ncRNA) also participates in the regulation of macrophage polarization [23,41–43]. At the present time, lncRNA is a hotspot in various research fields, and several reports have confirmed that lncRNA plays vital roles in macrophage differentiation and activation [10–14]. Our study was designed to identify novel and functional lncRNA molecules in two major patterns of macrophage activation.

In the present study, we analyzed lncRNA expression profiles in polarized BMDMs to uncover the potential role of lncRNAs in macrophage polarization via high-throughput microarray techniques. Among the thousands of identified differentially expressed lncRNAs, we found 627 that were up-regulated and 624 that were down-regulated in M(LPS+IFN-γ) macrophages compared to M(IL-4) macrophages. Following this, four lncRNAs (*AK048798, AW112010, 4933417E11Rik*, and *Dnmt3aos*) were selected for further confirmation by RT-qPCR. *AK048798* was selected because its potential target gene is Hif1α via trans-target prediction (data were not shown). This gene encodes a protein that has been shown to modulate macrophage polarization and to function in many diseases. *Dnmt3aos* was selected because it is a known lncRNA with no reported functionality. It locates on the antisense strand of Dnmt3a, which is an important component involved in epigenetic regulation [44,45]. The other two lncRNAs were randomly selected. The data showed that the RT-qPCR results were consistent with the microarray data.

Mechanistically, lncRNAs regulate gene expression via distinct processes that promote or inhibit recruitment of epigenetic regulators to modulate chromatin structure. Recent reports have shown that antisense lncRNA may function in sense-antisense pairs [28–31]. Our results showed the same regulation pattern. We saw that knockdown of *Dnmt3aos* expression altered the expression of *Dnmt3a* at both the mRNA and the protein levels in primary BMDMs, but silencing *Dnmt3a* did not alter the expression of *Dnmt3aos* (Fig 3). We also found that both the expression levels of *Dnmt3aos* and *Dnmt3a* were correlated to macrophage polarization. Meanwhile, we observed that *Dnmt3aos* was highly enriched in the nucleus, and *Dnmt3a*, as one of the DNA methyltransferases, catalyzed *de novo* DNA methylation in the nucleus. *Dnmt3a* plays an important role in cell development and tumorigenesis. DNA methylation represents one of the major epigenetic modifications and plays key roles in a variety of regulatory mechanisms of life processes [46–47]. The DNA methyltransferase (DNMT) family contains DNMT1, DNMT3A, and DNMT3B. DNA methylation in macrophage polarization is rarely reported [15] and lncRNA-mediated epigenetic modifications have not been previously examined. In our study, we determined that the DNA methylation levels in the two distinct kinds of polarized macrophages were significantly different. Notably, the pattern of DNA methylation also was found to be different in the two macrophage extremes. MeDIP-seq results showed that there were significant differences in genome-wide methylation levels between the si-NC and si-Dnmt3aos-transfected M(IL-4) macrophages. Combined analysis of DMR-related genes and mRNA expression profiles (reported in the literature 22) revealed a significant correlation between differential DMR-related genes and differential mRNA (Fig 5 and Fig 6). Our new results demonstrate that the *Dnmt3aos-Dnmt3a* interaction could be a key mechanism in macrophage polarization.

In fact, an altered genomic DNA methylation pattern characterized by regionally modified patterns of hypo- and hypermethylation exists in the si-NC and si-Dnmt3aos-transfected M(IL-4) macrophages. Notably, there was no major change in the methylation pattern at CpG islands, and the abnormal hypo- and hypermethylation status existed mostly in regions where CpG sites were sparse. Theoretically, reducing DNA methylation at gene promoters is associated with gene activation [48]. Our data also showed that inhibition of *Dnmt3aos* or *Dnmt3a* expression significantly increased the expression of the DMR enriched macrophage polarization related genes *IFN-γ, Cd24a, Icosl, Lox, TNF-α*, and *Lrrk2*. Then we confirmed the DNA methylation level by using BSP sequencing and bisulfite pyrosequencing. The results showed that among the randomly selected five TA clones used in the BSP DNA sequencing of *Lox* gene, the DNA methylation levels were hypomethylated in the si-NC group compared to the si-Dnmt3aos group (S5a and S5b Fig), and for *Icosl* gene, there was no difference between two groups (S5c and S5d Fig). The other four genes (*IFN-γ, Cd24a, TNF-α*, and *Lrrk2*) all contain the CG sites that were hypermethylated in the si-NC group compared to the si-Dnmt3aos group. We believe that BSP and pyrosequencing can also detect DNA hypomethylation and mixed methylation, and pyrosequencing can reliably detect DNA methylation differences among different cell populations, because bisulfite-treated DNA need not be cloned into bacterial expression vectors to avoid false positive results caused by random selection of clones. The corresponding selected hypermethylated CG sites in the si-NC group from the above four genes were validated through pyrosequencing. In summary, from the BSP and pyro-sequencing results, we find that only the CG dinucleotide locus at site 2 of the *IFN-γ* CG sites in the DMR region was confirmed to be significantly differentially methylated. The fact that the sequencing results were not consistent with the MeDIP-Seq results may possibly be because the MeDIP-seq enriched DNA sequences were sequence tandem repeats, which can be immunoprecipitated by 5-mC antibody, causing false positive results. The middle three CpG sites of the *IFN-γ* enriched DNA sequences could not be successfully sequenced for the same reason. On the other hand, as BSP sequencing required the cloning of bisulfite-treated DNA into bacterial expression vectors, and then randomly select positive clones for subsequent PCR and sequencing. However, the number of five clones selected by BSP sequencing in this study is relatively small and there is a great chance, so we think that BSP results may not represent the actual level of methylation. Bisulfite pyrosequencing, because of its own limitations on the length of sequencing fragments, is based on the results of BSP sequencing to select each gene difference CG site for pyrosequencing verification. The other three genes, *Cd24a, TNF-α*, and *Lrrk2* pyrosequencing showed no significant methylation differences at CG loci (S6-8 Fig), which may be due to the above reasons. Therefore, our data suggest that lncRNA *Dnmt3aos* regulates *Dnmt3a* expression leading to aberrant DNA methylation in macrophage polarization and *Dnmt3aos* specifically regulates the DNA methylation patterns of genes. In addition, mechanisms by which *IFN-γ* activates genes to promote macrophage activation via the Jak-STAT1 signaling pathway are well studied. A recent study has indicated that *IFN-γ* suppresses the basal expression of genes corresponding to an M2-like phenotype in *IFN-γ*-primed human macrophages [49]. In view of this, our study did not further explore the *IFN-γ* function in macrophage polarization.

In summary, this present study has determined the expression profiles of lncRNAs in macrophage polarization and contributes to the growing understanding of the role of lncRNAs in macrophage exposure to different stimuli. We found that *Dnmt3aos* plays a vital role in macrophage polarization. We also examined the global DNA methylation profiles between the si-NC and si-Dnmt3aos-transfected M(IL-4) macrophages, and demonstrated that *Dnmt3aos-Dnmt3a* axis-mediated aberrant DNA methylation may play a key role in regulating macrophage polarization. Our study enriches our understanding of the mechanism of macrophage polarization.

## Materials and Methods

### Ethics statement

All animal experimental procedures were approved by the Animal Ethics Committee of Wannan Medical College (Wuhu, China) and were performed according to the guidelines for the Care and Use of Laboratory Animals (Ministry of Health, China, 1998). The animals were sacrificed by cervical dislocation following anesthetized with a mixture of isoflurane and oxygen (3% v/v). All efforts were made to minimize the suffering of the animals.

### Animals

BALB/c mice 6-10 weeks of age and 25-30 g in weight were purchased from the Experimental Animal Center of Qinglongshan (Nanjing, China), were housed in pathogen-free mouse colonies, and fed a chow diet. Mice were kept in 12 - 12 hours of light-dark cycle; food and water were available ad libitum.

### Primary BMDM Isolation and Culture

Primary bone marrow-derived macrophages (BMDMs) of mice were obtained as described previously [22,23]. Briefly, BMDMs isolated from the femurs and tibias of mice were cultured with DMEM supplemented with 20% fetal bovine serum (FBS, Gibco), together with 20% L929 cell supernatant on 10 cm cell culture dishes at 37°C and 5% CO_2_. After 7 days of culture, the medium was removed, and the cells were cultured in fresh RPMI-1640 supplemented with 10% FBS for an additional 24 h. To induce the polarization of macrophages, BMDMs were then treated with DMEM/10% FBS containing 100 ng/ml LPS (Sigma) and 20 ng/ml IFN-γ (PeproTech) to produce M(LPS+IFN-γ) polarization, or with 20 ng/ml IL-4 (PeproTech) to produce M(IL-4) macrophages. After 48 h stimulation, the resulting cells were harvested for identification by morphology observation and flow cytometry (FCM) assay as in our previous publication [22–24].

### Microarray and Bioinformatic Analyses

An SBC mouse (4*180K) LncRNA microarray was designed for the global profiling of mouse LncRNAs and protein-coding transcripts. The sample preparation and microarray hybridization were performed as we have described previously [22]. Briefly, the total RNA from each sample was amplified and labeled by using a Low Input Quick Amp WT Labeling kit (Agilent Technologies). Labeled cRNA was purified using an RNeasy Mini kit (Qiagen GmbH). The concentration and specific activity of the labeled cRNAs (pmol Cy3/μg cRNA) were measured using a NanoDrop 2000. Each SBC mouse (4*180K) LncRNA microarray slide (Agilent Technologies Inc.) was hybridized with 1.65 μg Cy3-labeled cRNA using a gene expression hybridization kit (Agilent Technologies, Inc.). After 17 h of hybridization, the slides were scanned using an Agilent Microarray Scanner G2565C (Agilent Technologies, Inc.). Approximately 33,231 lncRNAs and 39,430 coding transcripts collected from the most authoritative databases, such as NCBI RefSeq, UCSC, ENSEMBL, FANTOM, UCR, and LNCRNA-DB were detected using microarrays. The Agilent Feature Extraction software (version 10.7; Agilent Technologies, Inc.) was used to analyze the acquired array images. Quantile normalization and subsequent data processing were performed using GeneSpring software version 11.0 (Agilent Technologies, Inc.). Microarray analysis was performed by Shanghai Biotechnology Corporation (Shanghai, China). Array data of differentially expressed protein-coding genes (fold change > 2 and *P* < 0.5) were deposited at the Gene Expression Omnibus database of the National Center for Biotechnology Information (accession no. GSE81922).

### Isolation of RNA and RT-qPCR

Total RNAs were extracted using the TRIzol reagent (Invitrogen). Cytosolic and nuclear fractions were prepared using the PARIS™ Kit (ThermoFisher Scientific) according to the instructions, and cDNA was synthesized using a SuperScript III First-Strand synthesis system (Life Technologies) with oligo(dT) and random primers according to the manufacturer’s instructions. Quantitative RT-qPCR was performed using QuantiTect SYBR^®^-Green PCR kits (Qiagen) and glyceraldehyde 3-phosphate dehydrogenase (*GAPDH*) or U6 as an internal control utilizing an Applied Biosystems Q3 real-time PCR system. The reactions were incubated in 96-well plates at 95°C for 3 min, followed by 40 cycles of 95°C for 15 sec and 60°C for 30 sec, followed by a dissociation curve. All the PCR reactions were run in triplicate. A complete list of primers used in this study is listed in Supplementary Table S1.

### RNA FISH

RNA FISH was performed as described by the manufacturer’s instructions as well as a previous publication [50]. In this study, a FAM-labeled locked nucleic acid (LNA) probe (Exiqon) against Dnmt3aos was used for RNA FISH. DAPI was used for nucleic counterstaining. Briefly, cultured cells were rinsed once with PBS and fixed with 4% paraformaldehyde (Sigma) in PBS for 20 min, then rinsed three times with PBS, permeabilized with 0.2% Triton X-100 (Sigma) in PBS for 15 min, washed twice with PBS, and incubated 30 min at 37°C with pre-hybridization buffer. Following this, cells were incubated 17 h at 37°C with hybridization buffer containing 500 nM FAM-labeled LNA probe. After successively washing with 4× SSC containing 0.5% Tween-20 three times, and 2× SSC, 1× SSC, and PBS once each, cells were stained with DAPI to visualize cell nuclei, rinsed three times with PBS, dried, and mounted in Vectashield mounting medium for fluorescent imaging. All immunofluorescence staining was photographed under either a confocal or an immunofluorescence microscope.

### Small Interfering RNA and Transfection of BMDMs

LncRNA smart silencer (a siRNA mix containing three ASOs and three siRNAs) targeting Dnmt3aos at different sites and small interfering RNAs (siRNAs) against Dnmt3a and negative control (NC) with no definite target were synthesized by RiboBio (Guangzhou, China). BMDMs were seeded on six-well plates at a density of 5 × 10^5^ cells/well overnight and then transfected with siRNA or the negative control at a final concentration of 100 nM using Lipofectamine 3000 (Invitrogen, USA). The interfering efficiency was detected by RT-qPCR or western blot 48 hours after transfection, and the siRNAs with silencing efficacy of more than 70% were selected for further experiments. Sequences of siRNAs are provided in Supplementary Table S2.

### Western blotting

BMDMs were transfected with either siRNA or the negative control as described above. At 48 h after transfection, cells were lysed in RIPA lysis buffer supplemented with cocktail protease inhibitor (Roche). Proteins were separated by SDS-PAGE and transferred onto a nitrocellulose filter membrane (Millipore, USA). The membranes were incubated with primary antibodies, either anti-Dnmt3a (1:1000, Abcam, USA), or anti-actin (1:1000, Santa Cruz, USA) in 5% milk/TBST buffer (25 mM Tris pH 7.4, 150 mM NaCl, 2.5 mM KCl, 0.1% Tween-20) overnight, followed by incubation with horseradish peroxidase (HRP)-conjugated anti-mouse or anti-rabbit IgG (Abcam) for 1 h. The target proteins were detected on the membranes by enhanced chemiluminescence (Millipore).

### Griess assay

Primary BMDMs were transfected with si-NC or si-Dnmt3aos for 48 h, followed by stimulation with LPS (100 ng/mL) plus IFN-γ (20 ng/ml) for an additional 6 h. The supernatants from the two groups of cells were collected, and then 50 μl aliquots of the conditioned medium were mixed with 50 μl of Griess reagent (Beyotime Biotechnology, China) and incubated for 10 min at room temperature in the dark. The colorimetric reaction was then measured at 540 nm using a Multiskan Go microplate reader (Thermo Scientific, Rockford, IL, USA).

### Arginase activity assay

Primary BMDMs were transfected with si-NC or si-Dnmt3aos for 48 h, followed by stimulation with IL-4 (20 ng/ml) for an additional 24 h. Briefly, 1 × 10^6^ treated macrophages were lysed with 50 μl of 0.1% Triton X-100 for 30 min and then added to 50 μl of 50 mM Tris-HCl/10 mM Cl_2_Mn·4H_2_O (pH 7.5) and incubated at 55°C for 10 min. L-arginine hydrolysis was carried out by incubating with 25 μl of 0.5 M L-arginine (pH 9.7) at 37°C for 60 min. The reaction was then stopped with 400 μl of stop solution (H_2_SO_4_ (96%)/H_3_PO_4_ (85%)/H_2_O (1:3:7, v/v/v)) and 25 μl of 9% of 2-isonitrosopropiophenone. The reactions were incubated at 100°C for 45 min and 100 μl of each sample was analyzed using a microplate reader at 540 nm. A standard curve was generated from urea solutions (0-20 mM), which were used to determine the final concentrations.

### Methylation Analysis by MeDIP Sequencing

DNA and RNA from the si-NC and si-Dnmt3aos-transfected M(IL-4) macrophages were extracted, and high throughput sequencing and MeDIP-seq were conducted by Shanghai Biotechnology Corporation, China. Detailed procedures were according to Chavez’s publication [51]. Briefly, DNA (3 μg) was sonicated at intensity 4 for 200 cycles per burst for 55 s (Covaris S2), and DNA fragments were end-repaired, ATP-tailed, and adapter-ligated with the Sample Preparation Kit (Illumina). Then DNA was recovered by AMPure XP Beads and used for methylated DNA immunoprecipitation (MeDIP) using the Magnetic Methylated DNA Immunoprecipitation Kit (Diagenode) according to the manufacturer’s protocol. After MeDIP, remaining DNA was PCR-amplified with sequencing primers and used for sequencing (Illumina Hiseq2500). Raw reads were preprocessed using the FASTX-Toolkit. MeDIP-seq peaks were counted using MACS. The depth of genomic regions was calculated and plotted using Integrative Genomics Viewer (IGV) and IGV Tools software.

### Bisulfite modification, bisulfite sequencing, and pyrosequencing

Genomic DNA was isolated from the si-NC and si-Dnmt3aos-transfected M(IL-4) macrophage cells and was then treated with bisulfite using the Imprint DNA Modification Kit (Sigma-Aldrich). Bisulfite sequencing and pyrosequencing were conducted by Shanghai Sangon Biotech Corporation, China. The procedure details were reported in previous publications [51,52]. Briefly, the bisulfite conversion-based PCR primers were designed with the MethPrimer program. Primers are listed in Supplementary Table S3. PCR was performed using P*fu* DNA polymerase (Sangon Biotech Corporation) under the following conditions: initial incubation at 98°C for 4 min, then 20 cycles of 94°C for 45 s, 66°C for 45 s, and 72°C for 1 min of touchdown PCR with a decrease of 0.5°C every cycle, continuing with 20 cycles of 94°C for 45 s, 56°C for 45 s, and 72°C for 1 min, followed by 72°C for 8 min for the gene promoter. The PCR products were purified using the Wizard DNA Clean-up System (Promega), and then cloned into the pGEM-T Easy Vector I (Promega). Five independent clones for each sample were picked, and the T7 and Sp6 primers were used to sequence inserted fragments. As for pyrosequencing, biotinylated DNA fragments (about 5 pmoles each) were attached to streptavidin-modified paramagnetic beads (Dynabeads™ M-280, Dynal A/S, Skøyen, Norway) by slow revolution of the suspensions in a thermostated (43° C) chamber (HB-1, Techne Ltd, Duxford, England) in a high salt (BW) buffer (0.05% Tween 20, 0.5 mm EDTA, 1 m NaCl, 5 mm Tris-HCl, pH 7.6). Following washing steps in BW buffer and 10 mm Tris-HAc, pH 7.6, the material was resuspended in 0.15 M NaOH. After a 10-min incubation period, the separated DNA strands were saved for subsequent processing as follows: beads were washed twice in 10 mm Tris-HAc, pH 7.6 and transferred to a solution (20 mm Mg(CH3COO)2, 10 mm Tris-HAc pH 7.6), containing 20-25 pmoles of the appropriate sequencing primer (primers are listed in Supplementary Table S4). Subsequently, the annealing procedure was conducted. This step comprised heating for 45 s at 95° C followed by cooling at room temperature. Throughout most of the preparation steps, the bead-coupled fragments were processed in parallel using a manifold device equipped with 96 magnetic ejectable microcylinders (PSQ 96 Sample Prep Tool, Pyrosequencing AB), which were protected by a disposable plastic pocketed cover. DNA sequence reading was conducted in a P3 PSQ 96 instrument (Pyrosequencing AB).

### Statistical Analyses

Descriptive statistics were generated for all quantitative data with presentation of means and S.D. Statistical analysis was carried out using Prism (GraphPad) software. Student’s *t* test was used to determine statistical significance, defined as *P* < 0.05.

## Acknowledgments

This project was supported by the National Natural Science Foundation of China (grant nos. 81472017, 81701557, 81802503, 81870017, and 81772180), Key Projects of Natural Science Research of Universities in Anhui Province (grant no. KJ2016A721 and KJ2018A0265). We thank LetPub (www.letpub.com) for its linguistic assistance during the preparation of this manuscript.

## Author Contributions

K. L. and Y. Z. designed the study. X. L. wrote the paper. M. Z. performed BMDM cells isolation and culture. H. Y. performed western blotting experiments. X. L. performed the other all experiments. X. K., W. P., Y. X., X. Z. and T. C. analyzed the data. M. Z. and J. Y. contributed reagents/materials and animal housekeeping. All authors approved the final version of the manuscript.

---

**S1 Fig.**
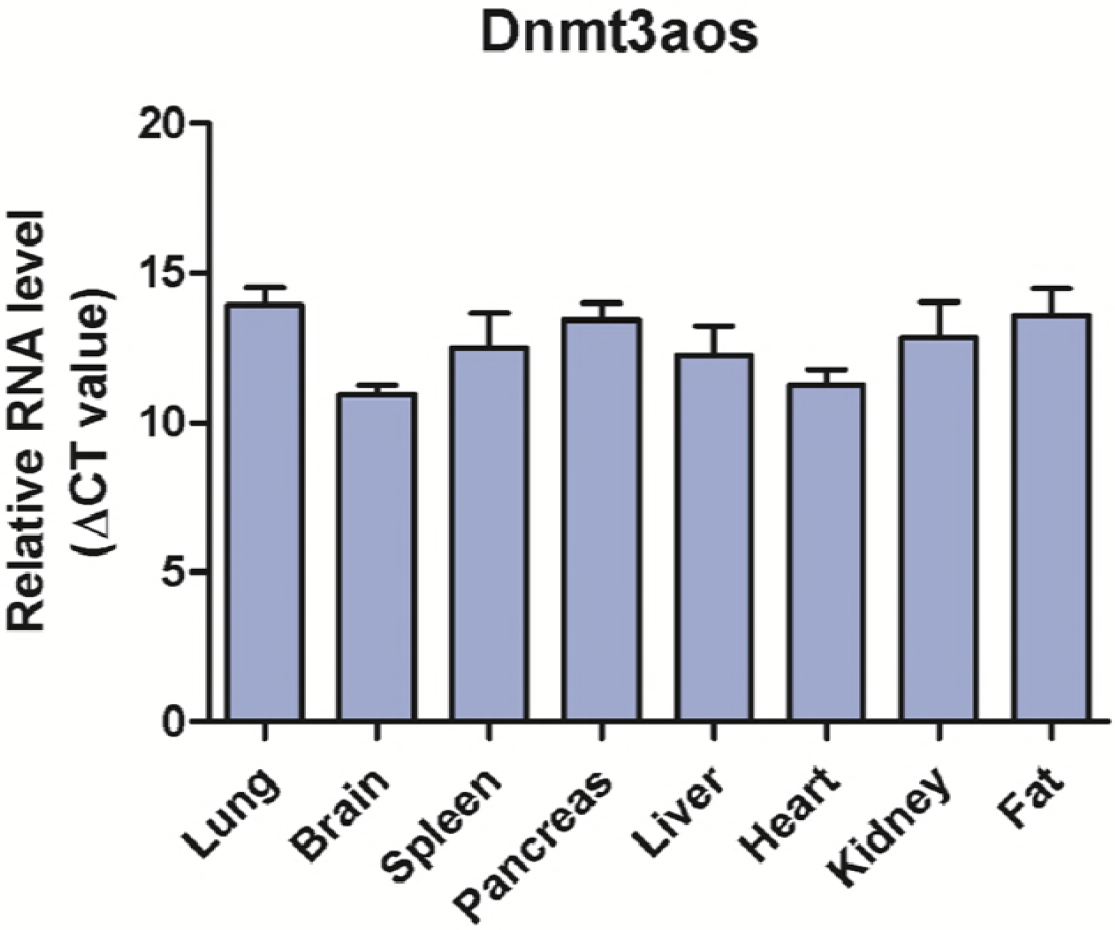
*Dnmt3aos* expression was detected in mouse tissues including lung, brain, spleen, pancreas, liver, heart, kidney, and fat. Error bars depict mean ± SD.

**S2 Fig.**
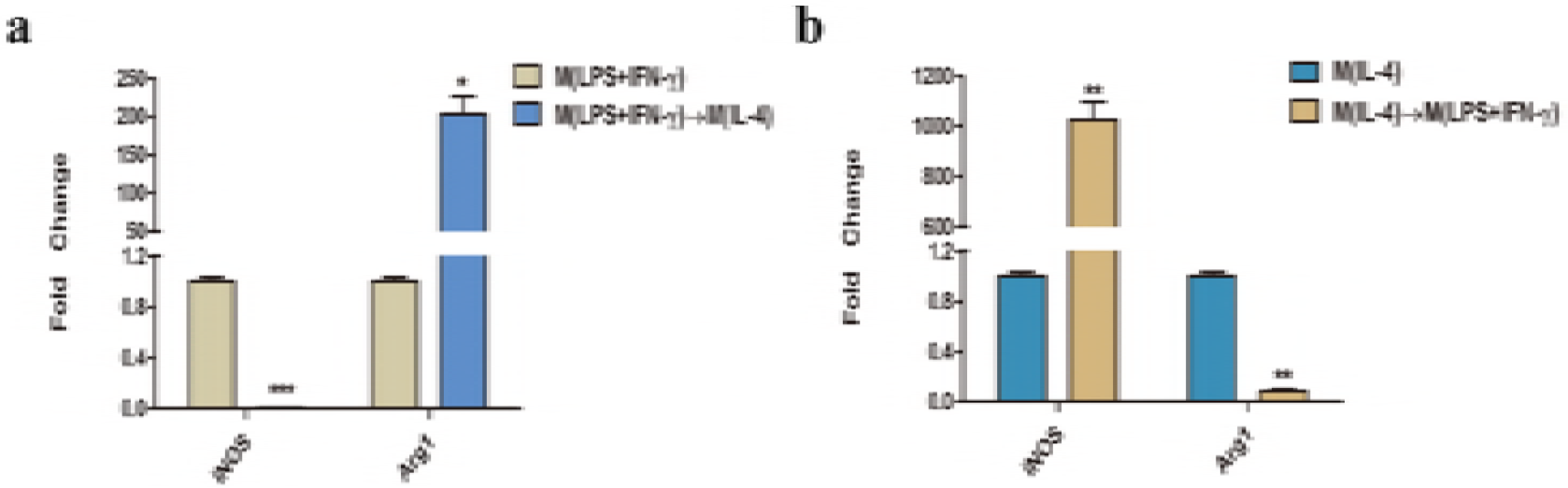
Relative gene expression during macrophage M(LPS+IFN-γ)/ M(IL-4) polarization. **(a)** Gene expression of iNOS or Arg1 following re-polarization of M(LPS+IFN-γ) macrophages to M(IL-4) macrophages by LPS/IFN-γ. **(b)** Gene expression of iNOS or Arg1 following re-polarization of M (IL-4) macrophages to M(LPS+IFN-γ) macrophages by LPS/IFN-γ. Data are representative of three separate experiments, and show the means ± SD, *P < 0.05, **P < 0.01, ***P < 0.001.

**S3 Fig.**
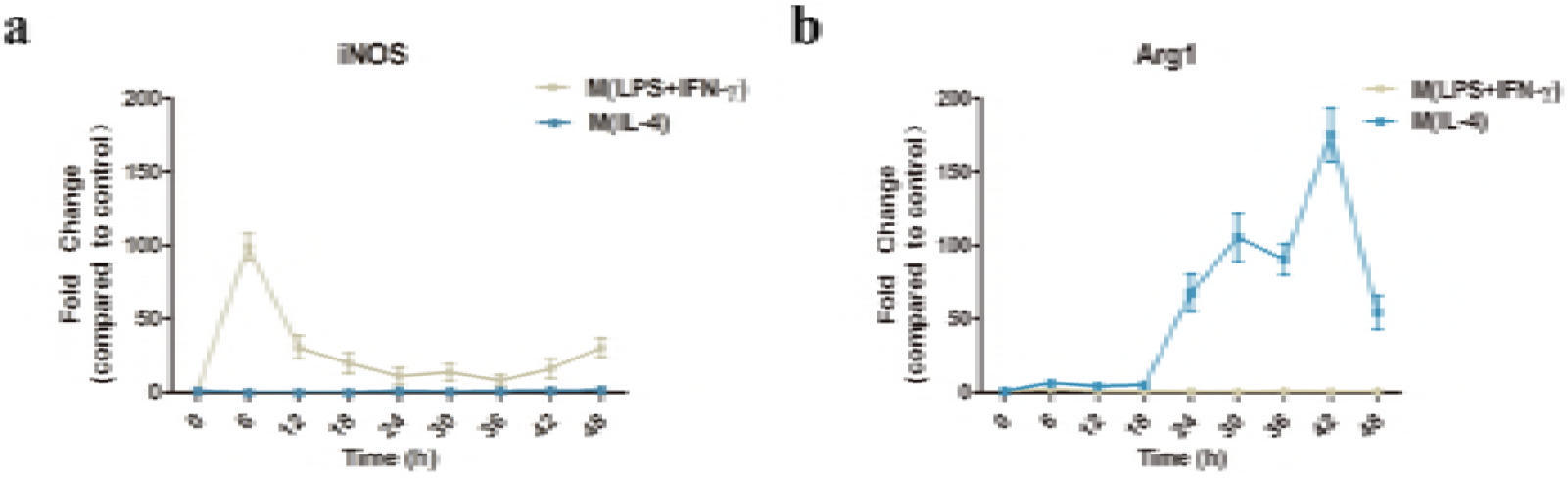
Identification of ex vivo-programmed M(LPS+IFN-γ) and M(IL-4) macrophages. BMDMs were cultured in the presence of LPS (100 ng/ml) plus IFN-γ (20 ng/m) or IL-4 (20 ng/ml). Polarization specific biomarkers were analyzed by RT-qPCR assays using RNA collected from BMDMs over different time courses post treatment. **(a)** iNOS **(b)** Arg1. Data are representative of three separate experiments and are compared to the control group.

**S4 Fig.**
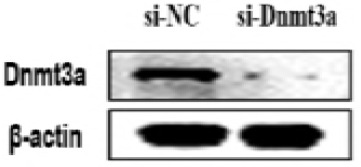
*Dnmt3a* siRNA inhibition efficiency in BMDMs was examined by western blotting.

**S5 Fig.**
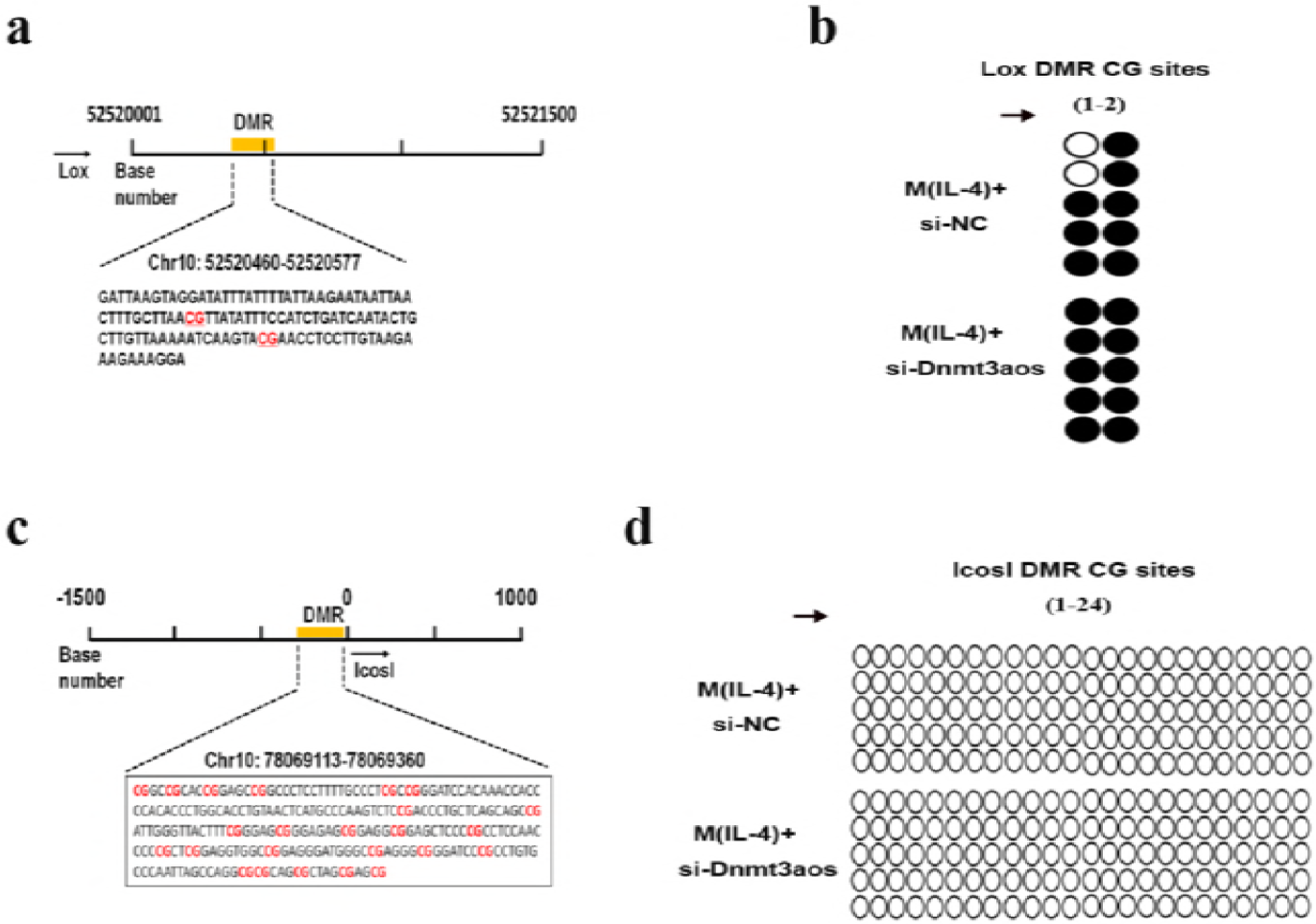
*Lox* and *Icosl* methylation profiles in the si-NC or si-Dnmt3aos transfected M(IL-4) macrophages. **(a)** The location and exhibition of DNA sequences of DMRs in the region of *Lox* obtained from MeDIP-seq enrichment. The CG sites (sites 1-2 in order) with underlining and red color were successfully BSP sequenced. **(b)** The open and filled circles symbolize the unmethylated and methylated CGs, respectively. Five colonies of separate CG sites from each group of *Lox* gene were further analyzed by BSP sequencing. **(c)** The location and exhibition of DNA sequences of DMRs in the region of *Icosl* obtained from MeDIP-seq enrichment. The CG sites (sites 1-24 in order) with underlining and red color were successfully BSP sequenced. **(d)** The open and filled circles symbolize the unmethylated and methylated CGs, respectively. Five colonies of separate CG sites from each group of *Icosl* gene were further analyzed by BSP sequencing.

**S6 Fig.**
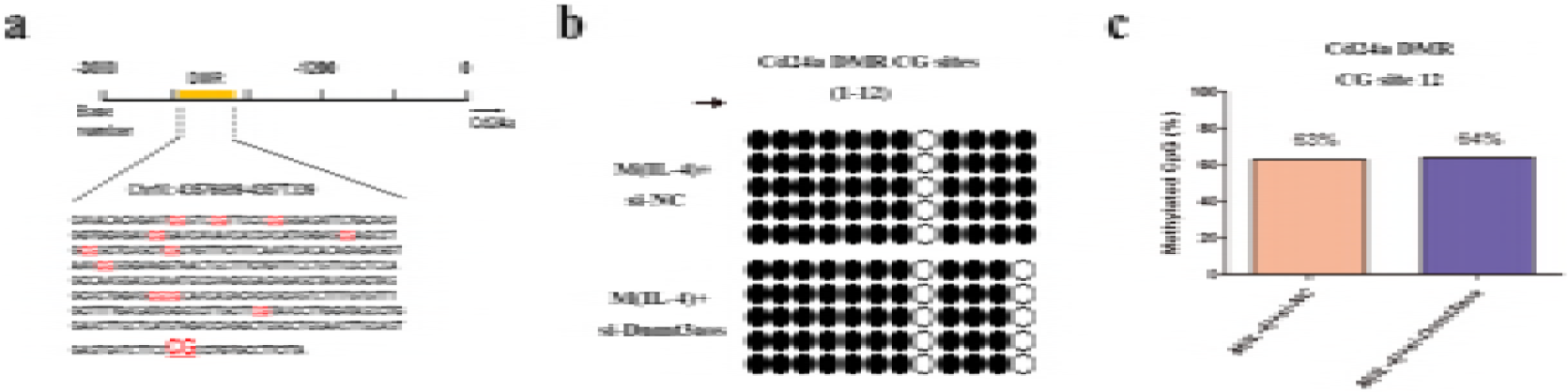
*Cd24a* methylation profiles in the si-NC or si-Dnmt3aos transfected M(IL-4) macrophages. **(a)** The location and exhibition of DNA sequences of DMRs in the promoter region of *Cd24a* obtained from MeDIP-seq enrichment. The CG sites (sites 1-12 in order) with underlining and red color were successfully BSP sequenced. **(b)** The open and filled circles symbolize the unmethylated and methylated CGs, respectively. Five colonies of separate CG sites from each group were further analyzed by BSP sequencing. **(c)** Pyrosequencing analysis of CG methylation level in the selected CG site of the *Cd24a* gene promoter from the BSP sequencing results. The figure illustrates the variation in methylation level at CG site 12 in si-NC or si-Dnmt3aos-transfected M(IL-4) macrophages.

**S7 Fig.**
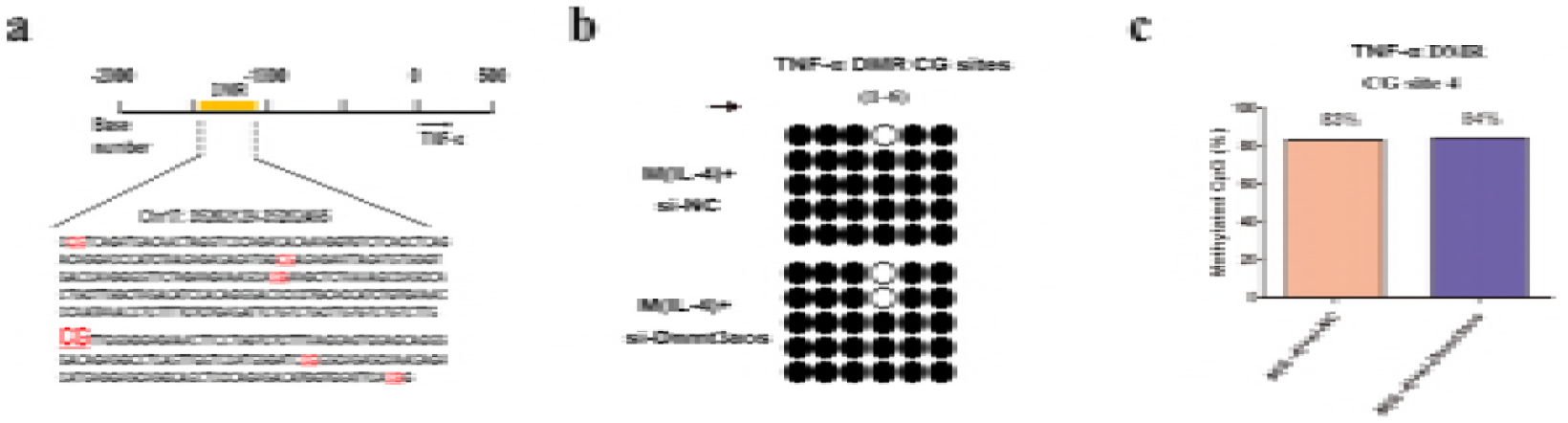
*TNF-α* methylation profiles in si-NC or si-Dnmt3aos-transfected M(IL-4) macrophages. **(a)** The location and exhibition of DNA sequences of DMRs in the promoter region of *TNF-α* obtained from MeDIP-seq enrichment. The CG sites (sites 1-6 in order) with underlining and red color were successfully BSP sequenced. **(b)** The open and filled circles symbolize the unmethylated and methylated CGs respectively. Five colonies of separate CG sites from each group were further analyzed by BSP sequencing. **(c)** Pyrosequencing analysis of CG methylation level in the selected CG site of the *TNF-α* gene promoter from the BSP sequencing results. The figure illustrates the variation in methylation levels at CG site 4 in si-NC or si-Dnmt3aos-transfected M(IL-4) macrophages.

**S8 Fig.**
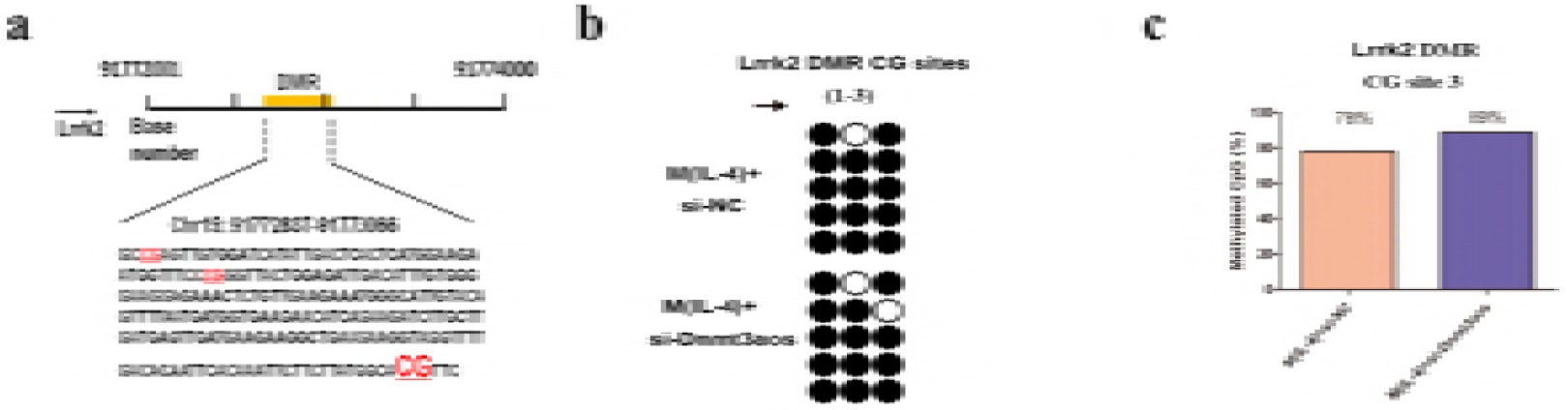
*Lrrk2* methylation profiles in si-NC or si-Dnmt3aos-transfected M(IL-4) macrophages. **(a)** The location and exhibition of DNA sequences of DMRs of *Lrrk2* obtained from MeDIP-seq enrichment. The CG sites (sites 1-3 in order) with underlining and red color were successfully BSP sequenced. **(b)** The open and filled circles symbolize the unmethylated and methylated CGs, respectively. Five colonies of separate CG sites from each group were further analyzed by BSP sequencing. **(c)** Pyrosequencing analysis of CG methylation levels in the selected CG site of the *Lrrk2* gene from the BSP sequencing results. The figure illustrates the variation in methylation levels at CG site 3 in si-NC or si-Dnmt3aos-transfected M(IL-4) macrophages.

**Supplementary Table S1.**
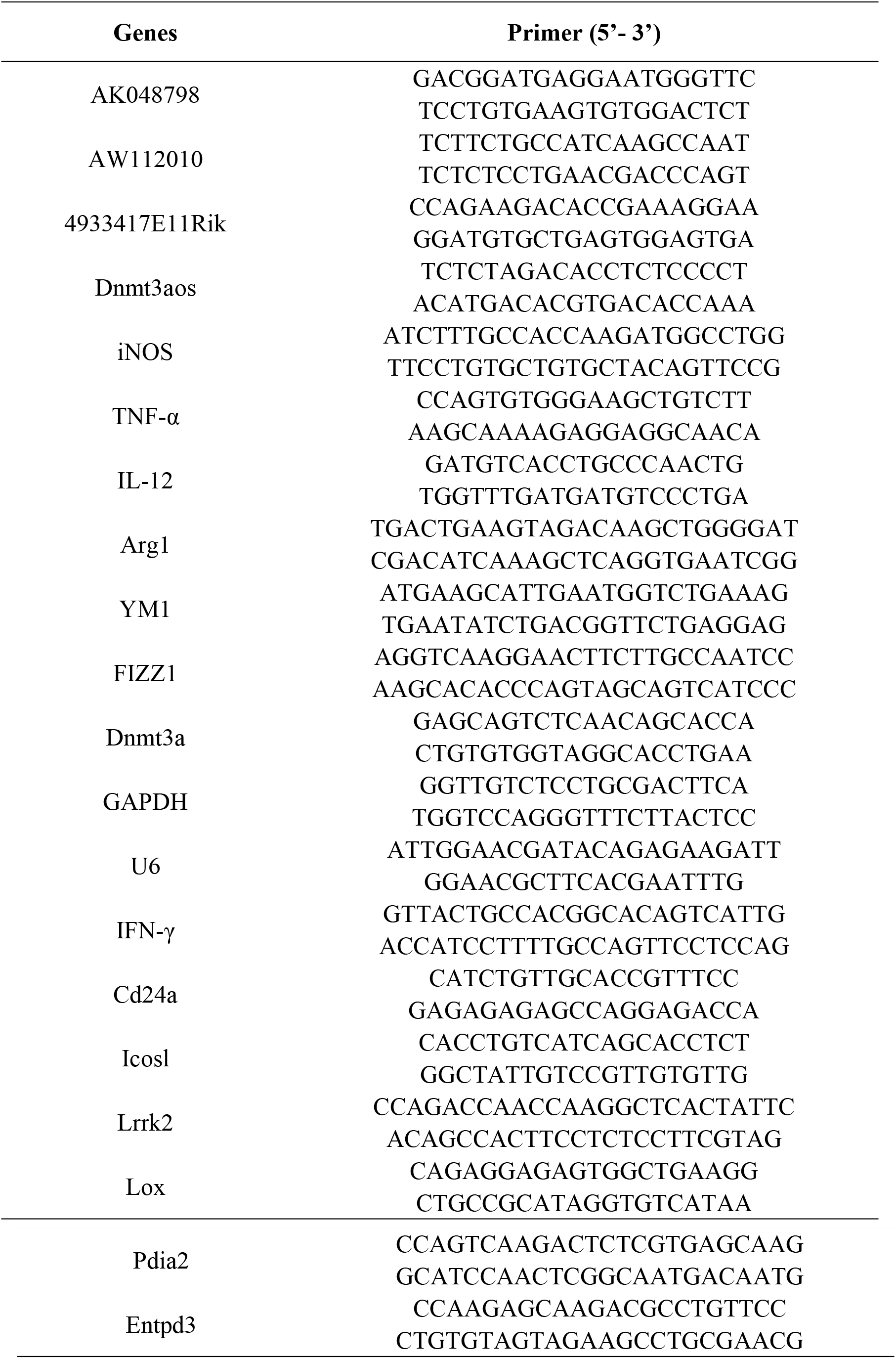
Primer sequences used in real-time PCR are written in 5’- 3’ direction.

**Supplementary Table S2.**
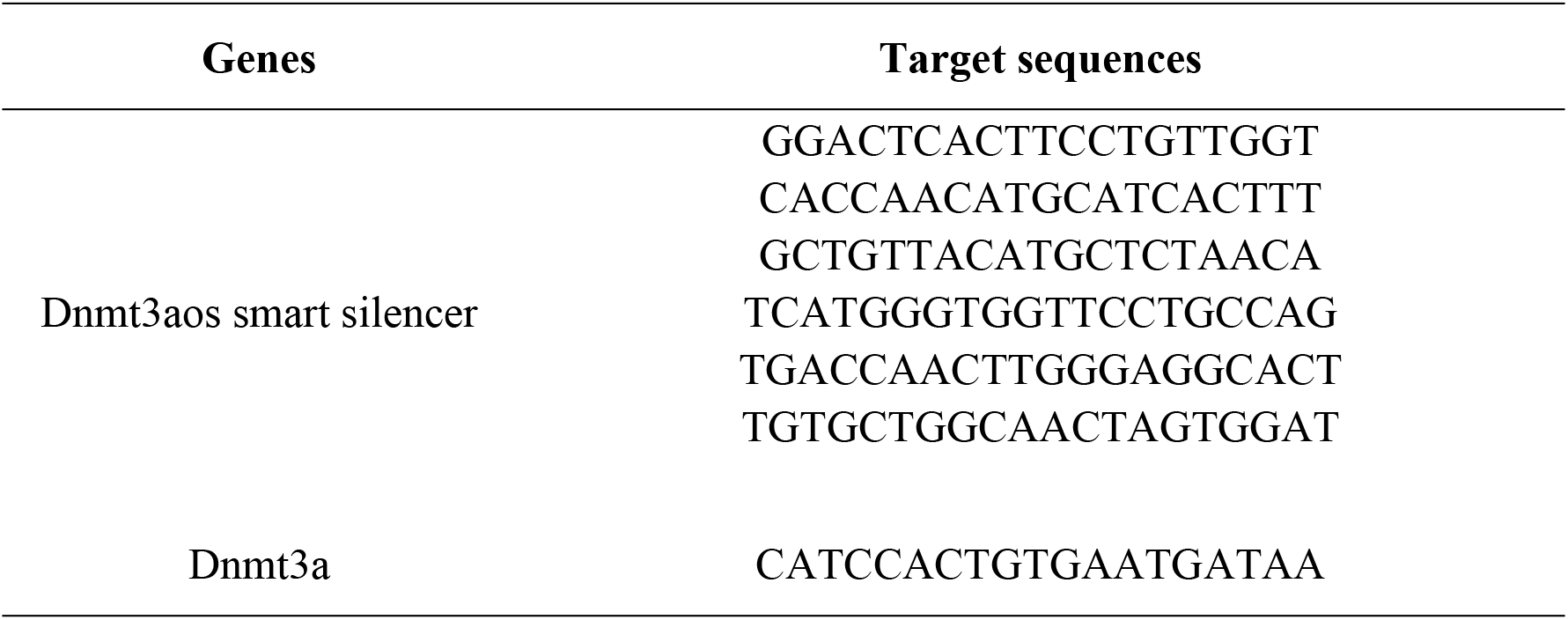
Oligonucleotides sequences used in siRNA transfection are written.

**Supplementary Table S3.**
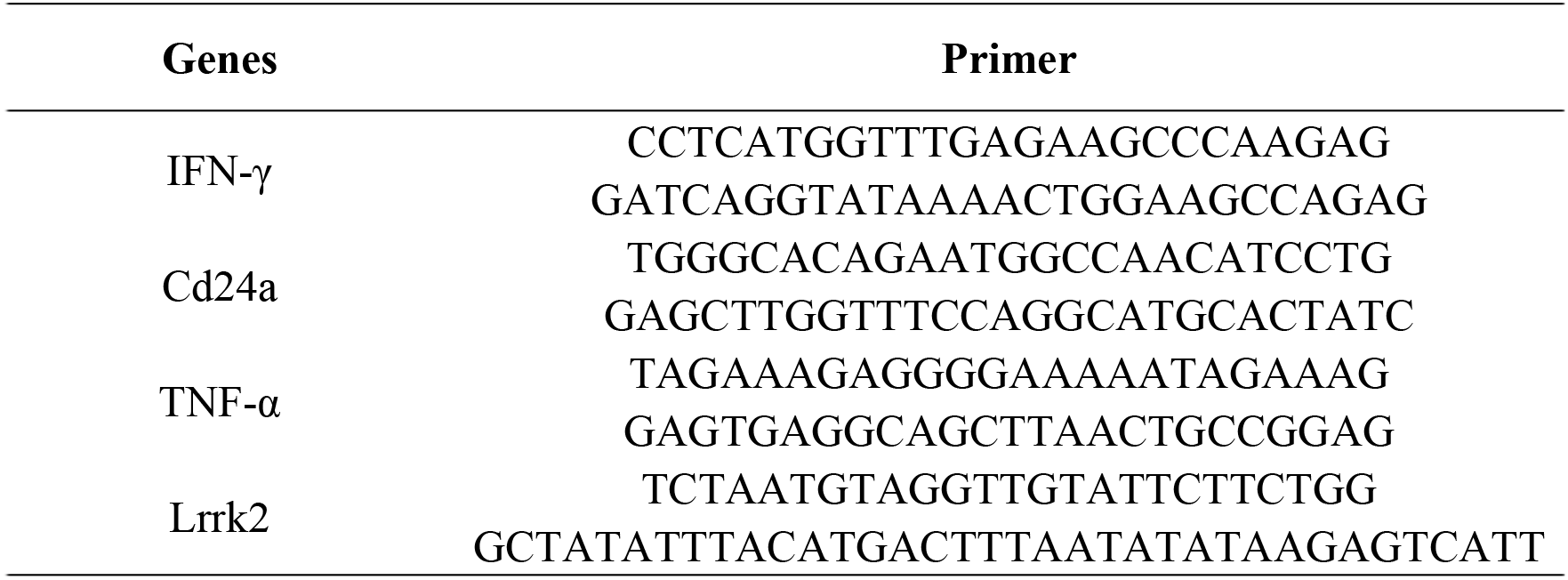
Primer sequences used in BSP sequencing are written.

**Supplementary Table S4.**
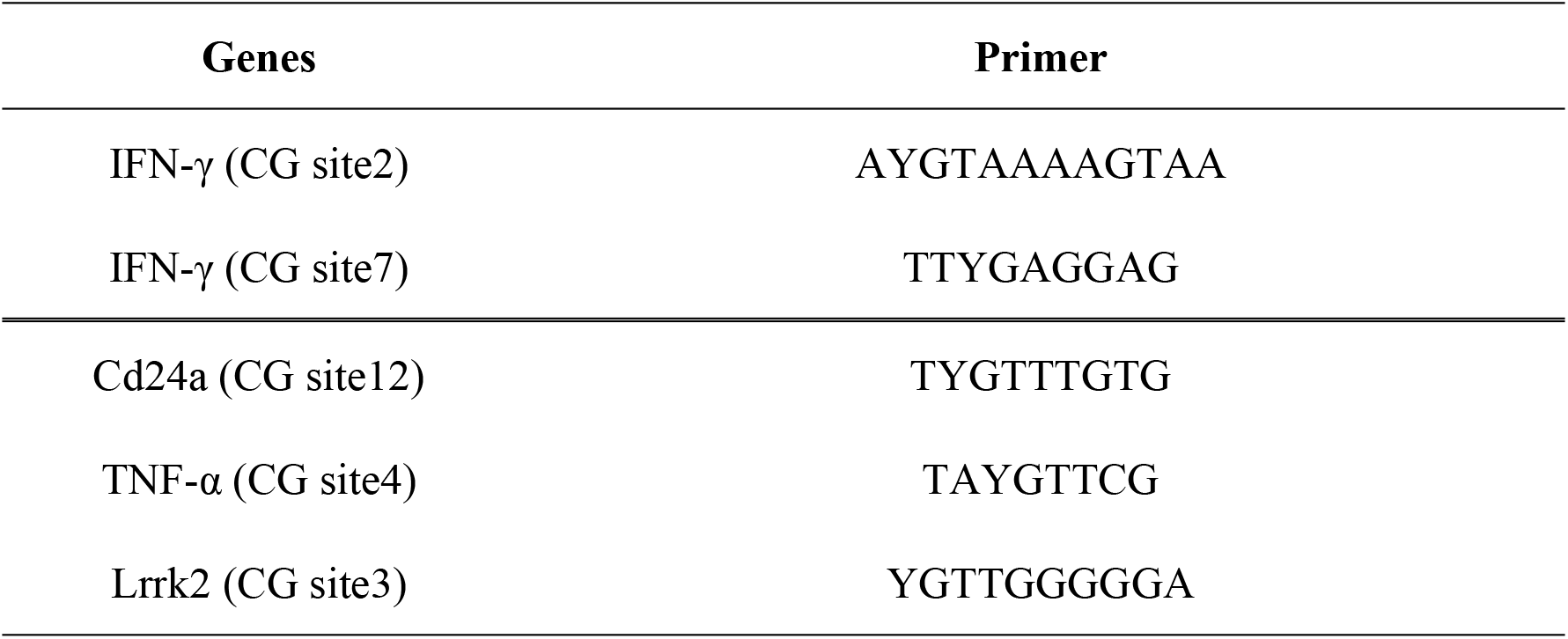
Primer sequences used in Pyrosequencing are written.

## Abbreviations

LncRNA: Long non-coding RNA
Dnmt3aos: DNA methyltransferase 3A opposite strand
Dnmt3a: DNA methyltransferase 3a
BMDM: bone marrow-derived macrophages
MeDIP-seq: methylated DNA immunoprecipitation sequencing
LPS: lipopolysaccharides
IFN-γ: interferon-gamma
GM-CSF: granulocyte-macrophage colony stimulating factor
IL-4: interleukin-4
IL-12: interleukin-12
IL-13: interleukin-13
TNF-α: tumor necrosis factor alpha
NO: nitric oxide
CD80: cluster of differentiation 80
CD68: cluster of differentiation 68
M-CSF: macrophage colony stimulating factors
TGF-β: transforming growth factor beta-1
Arg-1: Arginase 1
YM1: chitinase 3-like 3
Fizz1: Resistin-like-α
DMR: differentially methylated region
UTR: untranslated region
CDS: coding sequence
TTR: transcriptional termination region
GO: gene ontology
DEG: differentially expressed gene
5-Aza-CdR: 5-Aza-2’-deoxycytidine
BSP: bisulfite sequencing PCR
ncRNA: non-coding RNA.

